# A cell atlas of the developing human outflow tract of the heart and its adult aortic valve derivatives

**DOI:** 10.1101/2023.04.05.535627

**Authors:** Rotem Leshem, Syed Murtuza Baker, Joshua Mallen, Lu Wang, John Dark, Andrew D Sharrocks, Karen Piper Hanley, Neil A Hanley, Magnus Rattray, Simon D Bamforth, Nicoletta Bobola

## Abstract

The outflow tract (OFT) of the heart carries blood away from the heart into the great arteries. During embryogenesis, the OFT divides to form the aorta and pulmonary trunk, creating the double circulation present in mammals. Defects in this area account for one-third of all congenital heart defect cases. Here, we present comprehensive transcriptomic data on the developing OFT at two distinct timepoints (embryonic and fetal) and its adult derivatives, the aortic valves, and use spatial transcriptomics to define the distribution of cell populations. We uncover that distinctive embryonic signatures persist in adult cells and can be used as labels to retrospectively attribute relationships between cells separated by a large time scale. Single-cell regulatory network inference identifies GATA6, a transcription factor linked to common arterial trunk and bicuspid aortic valve, as a key regulator of valve precursor cells. Its downstream network reveals candidate drivers of human cardiac defects and illuminates the molecular mechanisms of both normal and pathological valve development. Our findings define the cellular and molecular signatures of the human OFT and its distinct cell lineages, which is critical for understanding congenital heart defects and developing cardiac tissue for regenerative medicine.

## Introduction

Congenital heart defects (CHD) are major birth abnormalities affecting ∼1% of new born babies ^1^. The outflow tract (OFT) carries blood away from the heart into the great arteries. During embryogenesis the OFT divides to provide separate aorta and pulmonary trunk vessels, which arise from the left and right ventricles respectively, giving the double circulation found in mammals ^2^. Defects specifically affecting the OFT of the heart represent a third of all CHD cases ^3^. In humans, the formation and remodelling of the OFT is a relatively rapid process, occurring in embryogenesis over a ∼ 4-week period from Carnegie Stage (CS)13 (4 weeks) to CS23 (8 weeks). Septation of the OFT begins around the CS14 stage when a transient aortopulmonary septal complex protrudes from the dorsal wall of the aortic sac ^4^. This divides the common OFT vessel into separate aorta and pulmonary trunks. The arterial (semilunar) valves are formed in the intermediate component of the OFT, initially as endocardial cushions ^5^ and continue to mature after septation of the arterial vessels. The morphological changes underlying OFT formation are orchestrated by two main cell lineages, the second heart field and the cardiac neural crest. The second heart field is a population of cardiac progenitor cells, originating from the pharyngeal mesoderm, that contribute to the formation of the myocardium of the right ventricle, atria and the OFT of the heart ^6^. The neural crest is a transient, pluripotent cell population, which migrate from the neural tube to multiple areas of the body. The cardiac neural crest, a sub-population of the neural crest, forms the aorticopulmonary septal complex, which separates the aorta and pulmonary trunk ^7–9^.

The use of model systems has substantially advanced our understanding of the cell lineages that contribute to the OFT. However, much less is known about OFT development in humans. In addition, relative to the embryonic period, adulthood is underexplored at the molecular level, and a clear relationship between cell lineages and cells found in the adult OFT is lacking. Previous studies have conducted single cell analysis of both developing and adult (whole or microdissected) hearts ^10–24^. Here, we present comprehensive transcriptomic data of the developing OFT (two distinct timepoints, embryonic and fetal) and its adult derivatives, the aortic valves, providing a large reference framework of OFT cell repertoires and their gene expression profiles. Using spatial transcriptomics, we describe the distribution of cell populations and cell–cell co-localizations. Remarkably, we identify the persistence of distinctive embryonic signatures in cells separated by a large timescale and use these signatures to establish lineage relationship between embryonic and adult cells. Our study defines the cellular and molecular signatures of the developing OFT and adult valves, and highlights the distinct cell lineages that construct these structures. This is of major importance for understanding the origin of congenital heart malformations and for producing cardiac tissue for use in regenerative medicine.

## Materials and methods

### Sample acquisition

Male CS16-17 (embryonic) and 12 pcw (fetal) OFTs were collected after pregnancy termination and snap frozen by the Human Developmental Biology Resource (www.hdbr.org). Embryonic samples were staged according to appearance by the HDBR embryo staging guidelines and pooled due to their limited size, with one CS16 and one CS17 sample in each pool. For fetal (12pcw) OFT, one OFT was used for each single nuclei RNA-seq, and one OFT for spatial transcriptomics. Adult aortic valve tissue was collected from three female adult (age 55-70) hearts declined for clinical transplantation with written informed consent for research from their families. Donors did not have ECHO evidence of aortic valve pathology or past medical history of aortic valve disease and inspection of aortic valve leaflets during dissection showed no signs of calcification. The research ethical approval (REC ref 16/NE/0230) was provided by the North East - Newcastle & North Tyneside Research Ethics Committee. The hearts were retrieved in the clinical standard fashion; arrested with 1 L of cold St Thomas’ cardioplegia solution at the agreed time point when both cardiothoracic and abdominal retrieval teams were ready for organ procurement. The donor hearts were then rapidly retrieved and preserved with either static cold storage or a hypothermic oxygenated perfusion device for 4 hours during which they were transported back to the laboratory at Newcastle University. After preservation, they were reanimated on a modified Langendorff system with blood-based oxygenated perfusate for 4 hours. At the end of the normothermic reperfusion, the donor hearts were dissected. Both left and right OFTs containing aortic and pulmonary valves respectively were excised, immediately snap frozen by being submerged into liquid nitrogen and stored at -80°C in The Newcastle Institute of Transplantation Tissue Biobank (17/NE/0022). The material was then obtained from The Newcastle Institute of Transplantation Tissue Biobank for analysis.

### Imaging

Human samples were processed for high-resolution episcopic microscopy and micro-computed tomography techniques as previously described. Briefly, aligned serial digital sections were imported into Amira (ThermoFisher Scientific) to produce two- and three-dimensional images ^25^.

### Nuclei extraction

Snap frozen embryonic and fetal tissue samples were minced using either a Dounce Homogeniser or a pellet pestle, then lysed for 30 min in ice cold lysis buffer (10 mM Tris HCl pH 7.4, 10 mM NaCl, 3 mM MgCl_2_, 0.1%Tween-20, 0.1% Igepal, 0.0005% Digitonin, 0.2 U/µl Protector RNAse inhibitor) or until no tissue pieces could be seen. Lysis was stopped with wash buffer (10 mM Tris HCl pH 7.4, 10 mM NaCl, 3 mM MgCl_2_, 0.1% Tween-20, 0.1% BSA, 0.2 U/µl Protector RNAse inhibitor) and filtered through a 20 µm filter. Nuclei were resuspended with PBS with Bovine Serum Albumin (0.1% BSA UltraPure^™^, AM2616 Invitrogen), and Protector RNAse inhibitor (0.2 U/µl, 3335399001 Roche), and processed using 10x Chromium Single Cell 3’ with target recovery set at 2500 nuclei/sample. Snap frozen adult aorta samples were cryosectioned to 50 µm, morphology was assessed by Hematoxylin and Eosin (H&E) staining and the valve tissue identified. Valve tissue from 3-4 unstained frozen sections was scraped off the slides, minced using pellet pestle and lysed in ice cold Igepal lysis buffer (10 mM Tris HCl pH 7.4, 10 mM NaCl, 3 mM MgCl_2_, 0.05% Igepal, 1 mM DTT, 1 U/µl Protector RNAse inhibitor) for 10 min. Sample was filtered using 40 µm cell strainer. After spinning down (500 rcf, 5 min, 4 °C) supernatant was removed and gently replaced with PBS/BSA (PBS containing BSA 1% and RNAse inhibitor 1 U/µl) and incubated for 5 min without disturbing the pellet. Supernatant was then removed, and the pellet resuspended in PBS/BSA supplemented with 7-aminoactinomycin D (7-AAD) dye. Nuclei were FACS sorted using a 100 µm diameter nozzle and a sheath pressure of 20PSI as per 10x protocols (document CG000375). 7AAD allowed nuclei to be identified from debris generated during the processing procedure. 7AAD stained nuclei were excited with a 488nm blue laser and emission was collected through a 685-725nm bandpass filter, at an event rate of ∼300-500 events per second. Following sorting, nuclei were permeabilized using 0.05X Lysis buffer (10 mM Tris HCl pH 7.4, 10 mM NaCl, 3 mM MgCl_2_, 0.01% Igepal, 0.001% Digitonin, 0.05 mM DTT, 1% BSA, 1 U/µl Protector RNAse inhibitor) for 1 minute, then washed and processed using 10x Chromium Single Cell 3’ with target recovery set at 2500 nuclei/sample.

### Visium spatial gene expression

One 12pcw whole fetal heart sample was embedded in OCT and cryosectioned at 5 µm according to 10x Genomics (document CG000240). A tissue optimisation was performed according to 10x Genomics protocol (document CG000238). Permeabilization time was set to 12 min. Four spatial gene expression sections were collected from the base of the aorta at regular intervals ending at the pulmonary valve. Tissue was processed and libraries created according to manufacturer instructions (document CG000239). 25% Ct value was determined by qPCR at 16 cycles.

### Single nuclei isolation and library construction

Gene expression libraries were prepared from single nuclei using the Chromium Controller and Single Cell 3ʹ Reagent Kits v3.1 (10x Genomics, Inc. Pleasanton, USA) according to the manufacturer’s protocol (document CG000315). Briefly, nanoliter-scale Gel Beads-in-emulsion (GEMs) were generated by combining barcoded Gel Beads, a master mix containing nuclei, and partitioning oil onto a Chromium chip. Nuclei were delivered at a limiting dilution, such that the majority (90-99%) of generated GEMs contained no nuclei, while the remainder largely contained a single nucleus. The Gel Beads were then dissolved, primers released, and any co-partitioned nuclei lysed. Primers containing an Illumina TruSeq Read 1 sequencing primer, a 16-nucleotide 10x Barcode, a 12-nucleotide unique molecular identifier (UMI) and a 30-nucleotide poly(dT) sequence were then mixed with the nuclear lysate and a master mix containing reverse transcription (RT) reagents. Incubation of the GEMs then yielded barcoded cDNA from poly-adenylated mRNA. Following incubation, GEMs were broken and pooled fractions recovered. First-strand cDNA was then purified from the post GEM-RT reaction mixture using silane magnetic beads and amplified via PCR to generate sufficient mass for library construction. Enzymatic fragmentation and size selection were then used to optimize the cDNA amplicon size. Illumina P5 & P7 sequences, i7 and i5 sample indexes, and TruSeq Read 2 sequence were added via end repair, A-tailing, adaptor ligation, and PCR to yield final Illumina-compatible sequencing libraries.

### Sequencing

The resulting sequencing libraries comprised standard Illumina paired-end constructs flanked with P5 and P7 sequences. The 16 bp 10x Barcode and 12 bp UMI were encoded in Read 1, while Read 2 was used to sequence the cDNA fragment. i7 and i5 sample indexes were incorporated as the sample index reads. Paired-end sequencing (28:90) was performed on the Illumina NextSeq500 platform using NextSeq 500/550 High Output v2.5 (150 Cycles) reagents. The .bcl sequence data were processed for QC purposes using bcl2fastq software (v. 2.20.0.422) and the resulting .fastq files assessed using FastQC (v. 0.11.3), FastqScreen (v. 0.14.0) and FastqStrand (v. 0.0.7) prior to pre-processing with the CellRanger pipeline.

## Data analysis

### Data pre-processing

Sequence files generated by the sequencer were processed using the 10x Genomics Cell Ranger pipeline. Developmental samples were analyzed with Cell Ranger v3.1.0, while adult samples used v6.1.2. FASTQ files were generated and aligned to a custom hg38 reference genome using default parameters. For developmental data, a custom premRNA_patched gtf file wsa used to include intronic reads, whereas for adult data the ‘include-introns’ option was enabled during the cellranger count runs. The pipeline identified cell barcodes corresponding to individual nuclei and quantified unique molecular identifiers (UMIs) per nucleus. Read alignment was performed using the STAR aligner, with multimapping reads excluded from UMI counting.

### Filtering

Low-quality nuclei were removed from the dataset using three commonly used parameter for cell quality evaluation, the number of UMIs per cell barcode (library size), the number of genes per cell barcode and the proportion of UMIs that are mapped to mitochondrial genes. The threshold of these three parameters were adjusted individually for each sample to retain only good quality nuclei from each of the samples. Outlier nuclei with a total read counts > 50,000 were removed as potential doublets. After filtering, the number of retained nuclei/total nuclei were as follows (all replicates are biological replicates): CS16-17_rep1 4703/7011; CS16-17_rep2 2576/7761; 12W_rep1, 10,151/12,110; 12W_rep2, 7306/16,110; AV1 2765/3008; AV2 446/609; AV3 2219/2362. To achieve a more fine-grained information for each dataset, each individual sample was first analyzed separately. Developmental time-points (embryonic and fetal) were then merged, and adult time-points were combined independently. Finally, all time points (developmental and adult) were integrated using scanpy ^26^, yielding a total of 30,166 nuclei for downstream analysis.

### Normalization and classification of cell-cycle phase

For individual sample analysis, raw gene expression counts were normalized using a deconvolution-based method ^27^. For integrated analysis in Scanpy, the default normalization approach was applied. Cell cycle phase scores (G1 and G2/M) for each nucleus were calculated using the cyclone method ^28^.

### Visualization & Clustering

The variance of each gene expression values was decomposed into technical and biological components; Highly Variable Genes (HVGs) were identified as genes for which biological components were significantly greater than zero. HVG genes were used to reduce the dimensions of the dataset using PCA; the dimension of dataset was further reduced to 2D using t-SNE and UMAP, where 1 to 14 components of the PCA were given as input. Nuclei were grouped into seven clusters using the dynamic tree cut method ^29^. For all datasets combined, scanpy’s highly_variable_genes (scanpy v. 1.9.5) were identified. Nuclei in scanpy aggregated data were clustered using Louvain clustering in scanpy with a resolution of 0.4.

### Identification of marker genes

For individual sample analysis, marker genes for each cluster were identified using the findMarkers function from scran 1.26.2 package ^30^. The function also returns FDR values for multiple testing correction using Benjamini-Hochberg procedure. Marker genes were then used to annotate the cell types of a cluster. For the scanpy aggregated data, scanpy’s rank_genes_groups was used to rank genes in each cluster using Wilcoxon rank-sum test method, which compares each cluster to the union of the rest of the clusters. This function also uses Benjamini-Hochberg procedure for multiple test correction. Gene ontology enrichment was evaluated for the top 100 genes using DAVID ^31, 32^, with P values limited to < 0.05.

### RNA-velocity

scVelo (v. 0.2.4) was applied to identify transient cellular dynamics in the embryonic stage ^33^. The spliced vs un-spliced ratio for each gene was calculated using the RNA-velocity’s command line tool, velocyto 10x with default parameters ^34^.

### SCENIC

The SCENIC pipeline was applied to infer gene regulatory network in our developmental samples. In the first step, GRNBoost2 created a gene co-expression module potentially regulated by the same TF, called regulon, followed by cisTarget to identify TFs that directly regulate each co-expression module, based on motif enrichments. Finally AUCell was used to calculate regulon activity scores in each cell. Output from AUCell was then used for downstream analysis using pySCENIC. Regulon Specificity Score (RSS) was used to quantify the activity of each regulon and identify cluster-specific regulons.

### Spatial Transcriptomics

Spaceranger v.1.3.1 was used to pre-process the Visium slides. After filtering the lower count spots and genes, data were normalized using scanpy’s normalize_total function. Spots were clustered using leiden clustering, and spatially variable genes were identified using Moran’s notes ^35^. As each of the Visium spots has more than one cell cell2location (v. 0.1.3) was used to deconvolute the spots ^36^. For cell2location genes were filtered using filter_genes() function and setting the parameter, cell_count_cutoff = 5, cell_percentage_cutoff2 = 0.03 and nonz_mean_cutoff = 1.12. The model was then trained using reference cell type signatures from snRNA-seq data, estimated using a negative binomial (NB) regression model. The Cell2location() function was then used to map the reference data to spatial spots while setting the N_cells_per_location parameter to 12 and detection_alpha to 20.

### Identification of gene expression patterns in spatial transcriptomics

To identify genes with expression patterns similar to a gene of interest, all spatial spots with expression values greater than 1 for the target gene were first selected. Genes not expressed in at least 50% of these spots were excluded. From the remaining set, any gene expressed in more than 25% of the spots where the target gene had zero expression was also removed. This filtering process yielded a list of genes with expression profiles closely matching that of the gene of interest.

### Lineage tracing algorithm

A transcriptional signature-based lineage inference method was developed to infer developmental relationships between embryonic populations and their fetal and adult counterparts. For each selected embryonic progenitor population (cluster), we defined a lineage signature by identifying the top 100 differentially expressed genes relative to other embryonic mesenchymal populations. In parallel, for each fetal or adult cell cluster, we identified the top 5000 differentially expressed genes by comparing expression within each cluster to all other clusters at the same (fetal or adult) stage. Lineage relationships were inferred by quantifying the overlap between each embryonic progenitor signature and the corresponding 5000-gene sets from later-stage clusters. Statistical significance of the observed overlaps was assessed using a hypergeometric test, enabling identification of fetal and adult populations significantly enriched for embryonic transcriptional signatures. Clusters exhibiting the strongest and most significant overlaps were interpreted as probable descendants of the corresponding embryonic progenitor populations.

All the experiments described (single nuclei RNA-seq and spatial transcriptomics) have been deposited in ArrayExpress; accession numbers are detailed in Table S1.

## Results

### Cellular landscape of the human OFT and its adult derivatives

To identify the cell types present in the human OFT, we conducted single nuclei (sn) RNA-seq on human OFT tissues. We isolated nuclei from 4 embryonic OFTs (CS 16-17), which were pooled into two independent pools (each from two embryos), two fetal (post conception week 12 (12pcw)) OFTs and from three adult aortic valves (Fig 1A; Table S1). The nuclei exhibit strong consistency between biological replicate samples (Fig 1B). After quality control and filtering, a total of 30,166 nuclei were segregated into 18 different clusters by unsupervised clustering (Fig 1C). These clusters were visualized by uniform manifold approximation and projection (UMAP). Each color on the chart represents a distinct cell population, arranged in order from the largest (5127 cells in cluster 0) to the smallest (355 cells in cluster 17). We decided against using batch-correction ^37^ to preserve the biological variability in our datasets, which reflects real changes across developmental time points (Fig S1AB). We observed the highest number of clusters (n = 12) in the fetal samples (Fig. 1D), consistent with new cell types arising from embryonic to fetal stage. The low number of clusters in the adult samples reflects sampling of the aortic valves, one of the derivatives of the entire OFT. Subsequently, we performed differential gene expression analysis to aid in the classification of each cell cluster (Table S2). Leveraging recognized markers, we were able to assign the clusters in developmental samples to three main compartments, cardiac, endothelial and mesenchymal cells (Fig 1EF). We also detected minor compartments of neuronal and immune cell types. In addition to mesenchymal (valve interstitial) and endothelial cell types, we allocated adult clusters to immune cells. We confirmed cell types in the main compartments using additional verified markers. The cardiac and endothelial nuclei populations appear to already express their appropriate cell type markers in the embryonic samples (Fig S1C). In contrast, most of the mesenchymal nuclei do not express markers of differentiated cell types (e.g. *DCN* or *MYH11*) at the earlier (embryonic) stage, but express a combination of *PDGFRA* ^38^ and *PDGFRB* ^39^, implying the embryonic stage captured mesenchymal nuclei before differentiation (Fig S1C). In addition, while embryonic and fetal cells display considerable variations in gene expression, adult cells share distinguishing features that set them apart from developing cells. This includes the expression of *NEAT1* ^40^, a long non-coding RNA (lncRNA), which does not distinguish cell types but discriminates adult from developmental tissues (Fig S1D-G).

**Figure 1.**
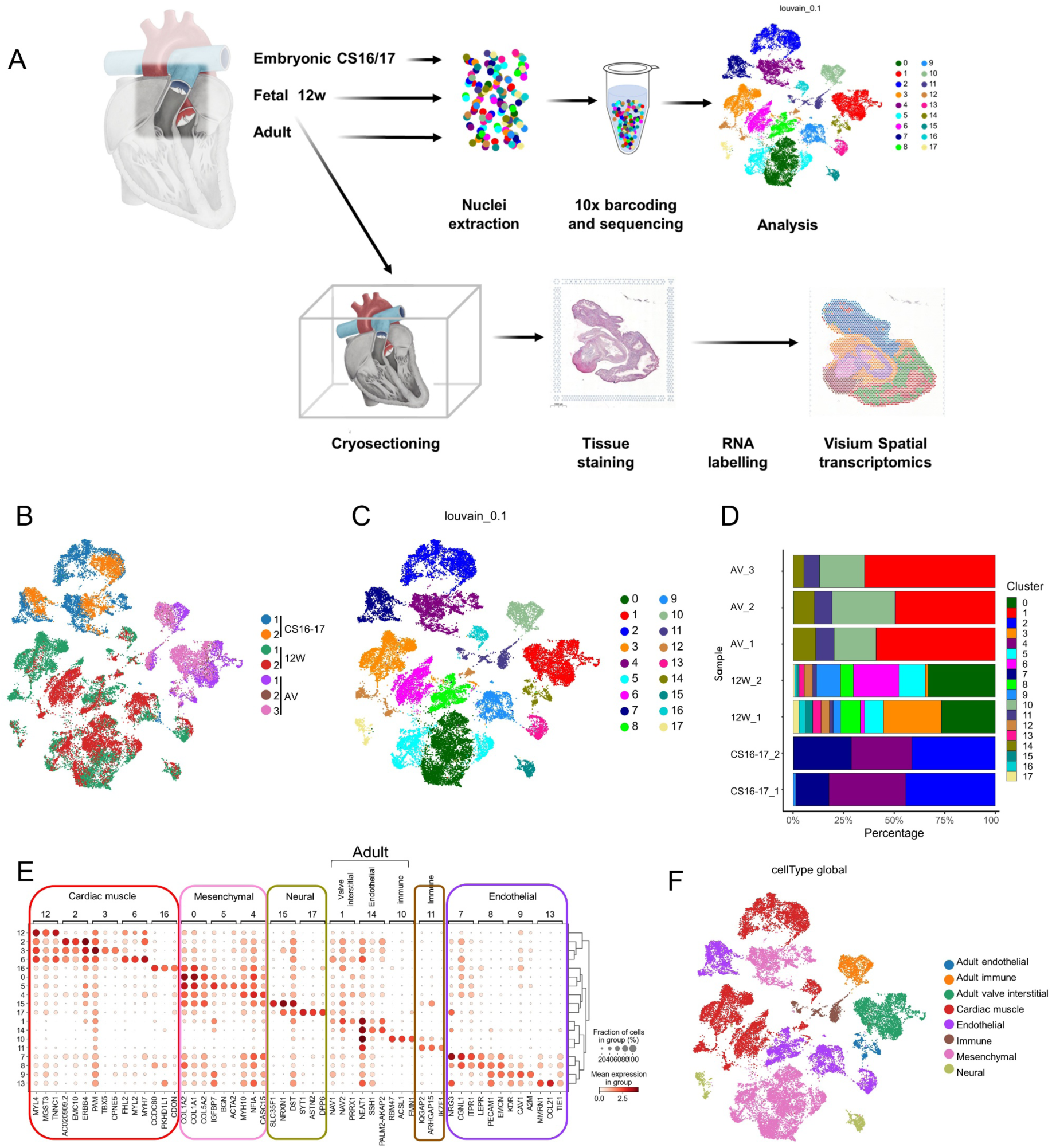
The cellular landscape of the developing OFT and its adult derivatives. A. Experimental schematics. Nuclei isolated from two embryonic (CS 16-17) and two fetal (12pcw) OFTs and from three adult aortic valves (AV) were analyzed by snRNA-seq. Four cryo-sections including the OFT region of a 12pcw heart were used in spatial transcriptomics (Visium). B. Sample correlation visualized by unsupervised clustering and projected on a two-dimensional UMAP. Nuclei are colored by sample, with embryonic (blue, orange), fetal (green, red) and adult (pink, purple and brown). C. Cell clusters visualized by the same UMAP as in B. Nuclei are colored by cluster. D. Cluster composition in each sample, presented as percentage of nuclei. E. Dotplot shows the mean expression levels of top differential genes across clusters and identifies five main cell types, cardiac, endothelial, mesenchymal (including valve interstitial), neural and immune. F. Cell types in E visualised by UMAP. Nuclei are colored by cell type. See also Fig S1.

### Identification of a GATA6-driven mesenchymal program underlying semilunar valve development

Given their critical role in OFT development, we focused our analysis on mesenchymal cells. To explore the heterogeneity within this lineage and uncover potential subtypes, we performed nuclei sub-clustering of embryonic and fetal datasets, followed by tSNE visualization (Fig S2AB). Using this approach, we obtained seven distinct mesenchymal clusters, two embryonic and five fetal (Fig 2AB). Fetal nuclei predominantly express *DCN*, a fibroblast marker, except for cluster 9, which contains *MYH11*-positive smooth muscle cells (Fig 2C, see also Fig S1C). In contrast, embryonic mesenchymal nuclei lacked definitive lineage markers, suggesting these cells are undifferentiated or at an early progenitor stage.

**Figure 2.**
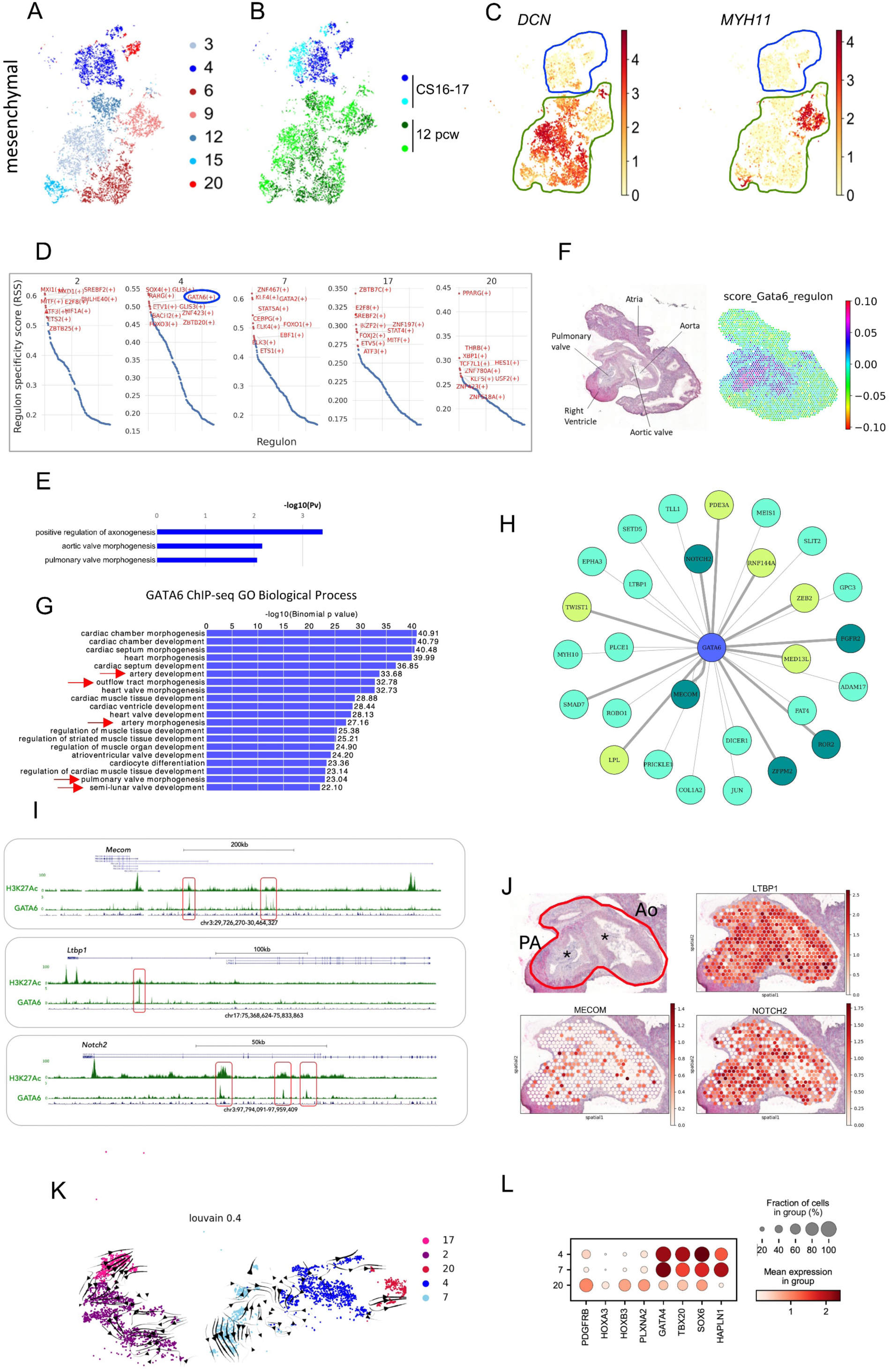
Characterization of embryonic mesenchymal nuclei. Mesenchymal cell clusters (A) and sample projection (B) of fetal and embryonic samples following subclustering, visualized on a two-dimensional tSNE. Nuclei are colored by cluster (A) and sample (B). C. Embryonic clusters (blue contour) do not express fibroblast (*DCN*) or smooth muscle (*MYH11*) markers, which are present in fetal nuclei (green contour). Nuclei are colored according to their scaled expression. D. Top 10 regulons in fetal clusters based on RSS score. E. Gene ontologies associated with the GATA6 regulon highlighted terms related to arterial and pulmonary valve morphogenesis. Functional annotation clustering of top 400 genes enriched GATA6 regulon was performed using DAVID and -Log10(Pv) was plotted in Excel. F. H&E-stained section showing the aortic and pulmonary arteries with their respective valves (left) and a corresponding map of the spatial expression of the GATA6 regulon (right). G. GREAT analysis of GATA6 high-confidence peaks (FE>10) in posterior pharyngeal arches and OFT at embryonic day (E) 11.5 (mouse). GATA6 peaks predominantly cluster around genes associated with cardiovascular terms, and specifically with OFT and artery development, as well as semilunar valve development (red arrows). H. Selected regulon genes associated with GATA6 binding in mouse embryo OFT and pharyngeal arches (see also Table S3) and associated with OFT-related abnormalities. Yellow genes are associated with human disease; light green genes cause mouse phenotypes; dark green genes are associated with both human and mouse defects. I. UCSC tracks of H3K27Ac ChIP-seq, GATA6 ChIP-seq (boxed in red) in posterior pharyngeal arches and OFT at E11.5 (mouse) and mammal sequence conservation at *MECOM* (top), *LTBP1* (middle) and *NOTCH2* (bottom) loci. J. Spatial Transcriptomics of Aorta (Ao) and Pulmonary Artery (PA) (clockwise): Hematoxylin and eosin (H&E) staining of the tissue area, with asterisks marking the semilunar valves; spatial distribution of *LTBP1, NOTCH2, MECOM*. K. Trajectory inference of future state of embryonic nuclei (CS16-17) showing mesenchymal (4, 20), endothelial-like (7) and cardiac (2, 17) clusters. Embryonic clusters derive from sub-clustering of aggregated fetal and embryonic nuclei shown in Fig S2A. L. Expression signatures in embryonic mesenchymal (4, 20) and endothelial-like (7) clusters. Both cluster 7 and 4 express high levels of cardiac TFs (*GATA4, TBX20*) and *HAPLN1*, a marker of semilunar valves. In contrast cluster 20 nuclei exhibit higher expression of neural crest markers, *HOXA3-B3* and *PLXNA2*.

Due to this absence of differentiated markers, which makes it challenging to define the identity of embryonic mesenchymal populations, we applied SCENIC ^41^ to the embryonic single-nucleus transcriptomes to infer key cell fate regulators and gain insight into their developmental potential. SCENIC links transcription factors (TFs) with their target genes based on co-expression, and it identified GATA6 as a top regulator of embryonic mesenchymal cluster 4 (Fig 2D). GATA6 is implicated in common arterial trunk (CAT) and bicuspid aortic valve (BAV) in humans ^42, 43^. Pathway enrichment analysis of the 400 genes in the GATA6 regulon highlighted GO terms related to arterial and pulmonary valve morphogenesis (Fig 2E), a process known to be regulated by GATA6, supporting the idea that this regulon contains functional downstream targets involved in valve formation. Consistent with this, spatial transcriptomic analysis of a later stage (12pcw) OFT shows that GATA6 regulon is mainly restricted to the aortic and pulmonary valves (Fig 2F). To prune TF-target interactions and identify GATA6 high-confidence direct targets, we used GATA6 genomic occupancy in the mouse OFT and posterior pharyngeal arches ^44^. Notably, GATA6 peaks are linked to GO terms highly specific to OFT and valve development ^45^, emphasizing that this dataset captures GATA6 binding to targets essential for the formation and maturation of these structures (Fig 2G). We inferred direct regulation by assigning peaks to genes ^45^. The GATA6 regulon is significantly enriched for genes occupied by GATA6 (p = 1.2 × 10⁻³³). This supports the interpretation that many genes within GATA6 regulon are associated with GATA6 binding, and likely to be direct GATA6 targets in the OFT. We systematically analysed direct targets for their implication in mouse phenotypes, human defects and genome-wide association studies (GWAS) using the MGI Database ^46^ and the EMBL GWAS database ^47^. The results are summarized in the network shown in Fig 2H. The GATA6 regulon includes genes identified in GWAS studies on aortic valve calcification (*RNF144A*, *ZEB2, MECOM, LPL, PDE3A, TWIST1*) ^48^ as well as genes whose inactivation in mice leads to phenotypes that overlap with GATA6 loss, including aortic valve and OFT defects (Table S3).

We detect strong GATA6 peaks overlapping with H3K27Ac, a histone mark associated with active enhancers, within the *Mecom* locus (Fig 2I). In mice, mutations in *Mecom*, which encodes a histone-lysine N-methyltransferase, result in CAT, interrupted aortic arch, and ventricular septal defects ^49^. In humans, GWAS have linked *MECOM* to calcified aortic stenosis ^48, 50^. *MECOM* transcripts are sparsely expressed, and primarily localized within the aorta and pulmonary artery, in regions occupied by the semilunar valves (Fig 2J). The *Notch2* locus also exhibits multiple GATA6 peaks in regions of high acetylation and evolutionary conservation across mammals (Fig 2I). Conditional deletion of *Notch2* in cardiac neural crest cells leads to hypoplastic aortas and pulmonary arteries due to reduced smooth muscle content ^51^. *NOTCH2* is one of the causative genes in Alagille syndrome, a multisystem disorder involving hepatic and cardiac anomalies, most commonly peripheral pulmonary stenosis and tetralogy of Fallot ^52^. Like *MECOM, NOTCH2* is primarily expressed in the aorta and pulmonary artery (Fig 2J). Strong GATA6 binding is also observed at the *Ltbp1* gene, which encodes an extracellular regulator of TGF-β signalling (Fig 2I). Loss of the long isoform of *Ltbp1* in mice results in CAT and interrupted aortic arch, due to defective cardiac neural crest cell function ^53^. Unlike *MECOM* and *NOTCH*2, *LTBP1* is highly expressed in the walls of the aorta and pulmonary artery, including the OFT septum (Fig 2J). Additional genes in the GATA6-regulated network include *SLIT2* and *ROBO1,* components of the SLIT-ROBO signaling pathway implicated in BAV ^54^, and structural genes like *MYH10* and *COL1A2* (Fig S2C). Collectively, the convergence of OFT phenotypes, OFT-specific GATA6 binding and enhancer activity (H3K27Ac enrichment), and the expression patterns of regulon genes supports a GATA6-driven transcriptional network implicated in OFT development, particularly the semilunar valves. These findings delineate molecular mechanisms underlying GATA6 function in the developing valves and highlight candidate genes that may contribute to BAV susceptibility.

SCENIC analysis links mesenchymal cluster 4 to semilunar valve development. We next applied trajectory inference to examine its relationship to other embryonic clusters at the same stage. RNA velocity predicts the future state of individual cells by distinguishing between un-spliced and spliced mRNAs ^34^. We detected the transition of some nuclei from cluster 7 (endothelial-like cells) specifically towards cluster 4 (mesenchymal cells) (Fig 2K). Cluster 7 exhibits a similar expression profile to cluster 4, but distinct from the other mesenchymal nuclei (cluster 20) (Fig 2L). This profile includes high expression of *HAPLN1*, encoding the extracellular matrix cross-linking protein Hyaluronan and Proteoglycan Link Protein 1, found in the endocardial lining and the developing semilunar valves in the OFT. Compared with fetal endothelial clusters, endothelial cluster 7 is enriched for genes associated with aortic and pulmonary valve morphogenesis and epithelial to mesenchymal transition (EMT) (Fig S2F; Table S4). In mice, semilunar valve formation begins with an EMT of endothelial cells in the endocardium, the cell layer adjacent to the myocardium^55, 56^. This suggests that cluster 7 likely contains an endocardial population, with RNA velocity tracking their transition into mesenchymal cells in cluster 4, thereby generating the precursors of valve interstitial cells.

To understand why GATA6 emerges as a top regulator specifically in cluster 4, we examined GATA6 expression across embryonic nuclei. Although *GATA6* is expressed in all embryonic clusters, its levels are highest in cluster 4 (Fig S2D), which may account for its restricted activity in this population. Alternatively, given that TFs typically act cooperatively, GATA6 may coregulate cluster 4 regulon in combination with additional factors.

To identify these additional factors, we compared the regulons of cluster 4 top transcriptional regulators (Fig 2D). As expected, since regulon genes are sampled from cluster 4 enriched transcripts, these regulators share many downstream targets with GATA6 (19-30% overlap). Notably, GLI3 shows substantially greater regulon overlap (56%), suggesting functional cooperation with GATA6. This is consistent with their reported cooperation in the developing mouse limb^57^. GLI3 is also enriched in cluster 4, further supporting the hypothesis that these TFs cooperate in the development and differentiation of this cell population (Fig S2E).

### Spatial resolution of mesenchymal nuclei in the OFT

Mesenchymal cells build key structures in the OFT, specifically the separation of the aorta from the pulmonary artery (Fig 3A, C, E) and the semilunar valves at the base of the aorta and pulmonary artery (Fig 3B, D, F). At 12pcw, mesenchymal nuclei express markers of differentiated cell types (Fig 2C). To visualize the distribution of fetal mesenchymal cell populations within the OFT, we conducted spatial gene expression analysis. We generated four sections of a 12pcw OFT (equivalent stage to snRNA-seq), starting at the level of the aortic valves (a) and ending at the level of the pulmonary valves (d) (Fig. 3G). We used Cell2location ^36^ to transfer the labels from the transcriptomic data to spatial gene expression data and mapped different cell-types on each individual spatial location. Cardiac nuclei correctly mapped to the atria and the ventricle (Fig S3A). Mesenchymal cells largely distributed within and around the vessels, and also mapped to the valves of the pulmonary artery and the aorta (Fig S3B). Endothelial cells were primarily located in the aortic valves (Fig S3C). Immune cells were scattered around the tissue (Fig S3D), while neural cells were concentrated in a spot of the atria (Fig S3E). Next, we mapped the five fetal mesenchymal clusters to distinct structures in the OFT (Fig 3H) and used distinctive markers to confirm spatial assignments. Clusters 3 and 6 (Fig 3I-I^I^) map to the arterial outer walls and the septum between the aorta and pulmonary arteries and largely colocalise with *DCN*-positive cells (Fig 3I^II^). These two clusters largely overlap. Cluster 9 (Fig 3J) coincides with *MYH11* expression (Fig 2J^I^): *MYH11* is a terminal marker of smooth muscle differentiation ^58^, which identifies the aortic smooth muscle layer. Cluster 12 (Fig 3K) concentrates to both aortic and pulmonary valves and co-localizes with *HAPLN1* (Fig 3K^I^), a marker of the semilunar valves ^59^. No molecular differences or distinguishing markers were identified between the aortic and pulmonary valves. Finally, the smallest cluster (15) is restricted to the cardiac ventricle and is *DCN*-negative, therefore was eliminated from our further analysis (Fig S3F). Thus, the five subtypes of mesenchymal nuclei, identified by snRNA-seq, largely correspond to spatially segregated cell populations of fibroblasts (cluster 3, 6), smooth muscle cells (cluster 9) and valvular interstitial cells (cluster 12). To facilitate accessibility to these data, we have created a cellxgene VIP app to visualize gene expression across four sections of the fetal heart, available at: https://cellxgene.cziscience.com/collections/5d2077ea-7b49-45c8-b4cb-64790b698591.

**Figure 3.**
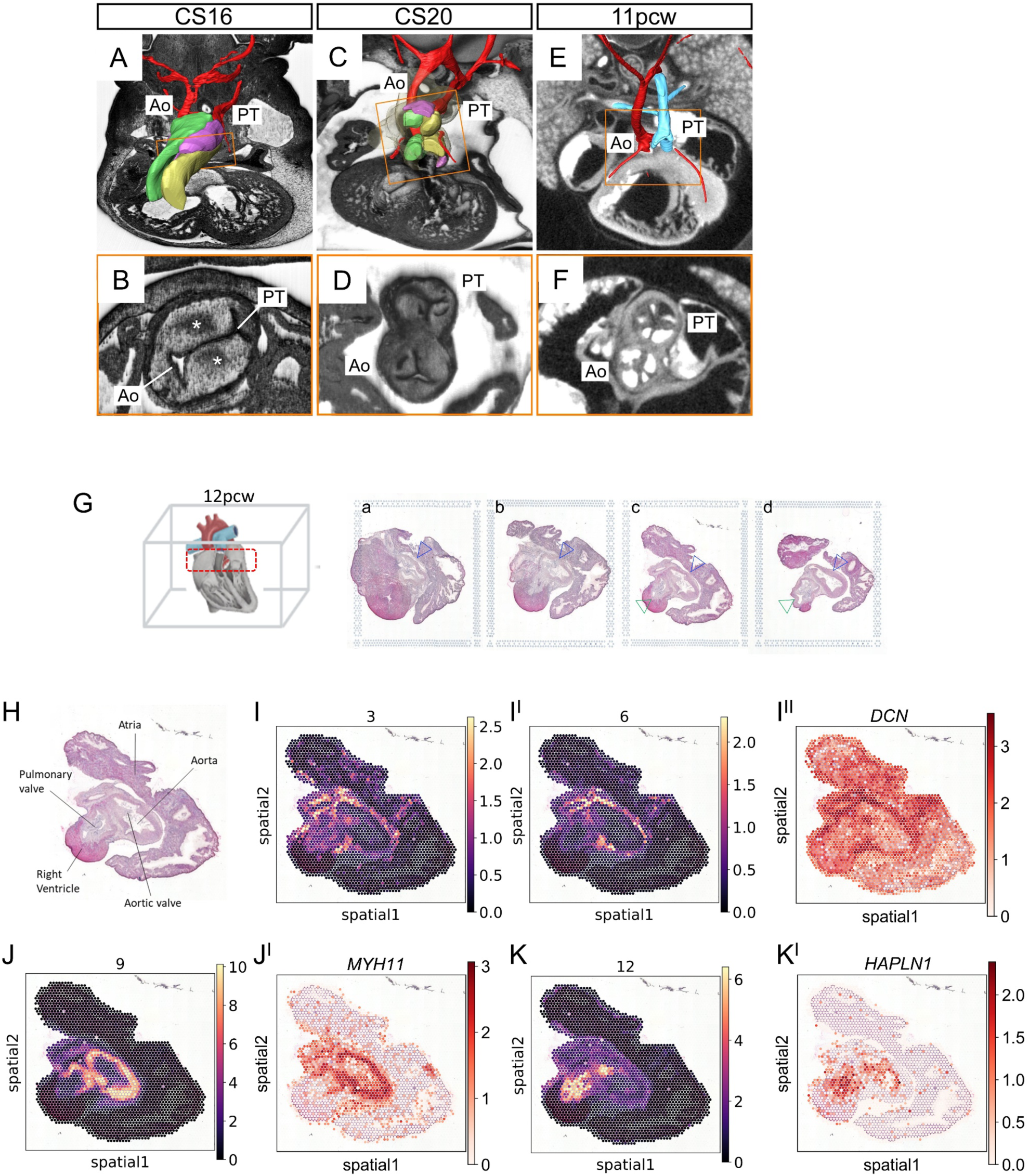
Spatial distribution of mesenchymal clusters. A-F. OFT valve formation and remodelling. Images were prepared using high resolution episcopic microscopy at embryonic stages (A-D) and by micro-CT at the fetal stage (EF). A. By CS16 the OFT has septated into the aorta (Ao) and pulmonary trunk (PT). The immature OFT cushions are visible: the septal (yellow), parietal (green) and intercalated (purple) cushions. B. Neural crest cells (asterisks) contribute to valve formation. CD. At CS20 the cushions have begun to remodel to form the three leaflets of the aortic and pulmonary semilunar valves. EF. At 11pcw the valves have transformed into the leaflets that control the unidirectional flow of blood from the heart. Boxed regions in A-C-E are shown at higher magnification in B-D-F. G. Heart alignment for sectioning, with the OFT region marked in red (left). H&E staining of OFT cryo-sections used for spatial transcriptomics from the base of the OFT (a) to the pulmonary valves (d). The aorta and pulmonary trunk are indicated by blue and green arrowheads, respectively. H. H&E section ‘c’ annotated to show major structures. I, I^I^, J, K. Spatial distribution of mesenchymal clusters (purple and yellow). I^II^, J^I^, K^I^. Spatial distribution of lineage specific markers (white and red). I-I^II^. Clusters 3 (I) and 6 (I^I^) largely overlap with fibroblast lineage marker *DCN* (I^II^). J-J^I^. Cluster 9 (J) and smooth muscle lineage specific marker *MYH11* (J^I^) map to the aortic walls as well as to the pulmonary artery. K-K^I^ Cluster 12 (K) and valve specific marker *HAPLN1* (K^I^) are mainly found in the valves at the base of the aorta and pulmonary artery. See also Fig S3.

### Developmental trajectories of mesenchymal cells in the developing OFT

At CS16-17, we identified two clusters of undifferentiated mesenchymal cells. By 12pcw, three main types of differentiated mesenchymal populations had emerged: fibroblasts, smooth muscle cells, and valvular interstitial cells, each occupying distinct spatial locations within the OFT (Fig 2A-B). Our objective was to track the developmental trajectories of these populations and identify the 12pcw descendant cells from each embryonic cluster. Connecting mesenchymal embryonic progenitors to their differentiated fetal counterparts is challenging, because the embryonic nuclei are yet to express the molecular markers of differentiated lineage descendants (Fig 2C). Trajectory inference methods ^60^ failed to establish lineage relationships between embryonic and fetal populations. To overcome this, we used gene module scores. The rationale behind this approach was that any distinctive developmental signature present in the embryonic clusters would likely be retained in the fetal nuclei, thereby enabling us to trace the trajectories of mesenchymal cell populations.

We first performed pairwise differential gene expression analysis of embryonic clusters to identify distinct developmental signatures of mesenchymal subtypes. Cluster 4, which partly derives from endocardial cells, is enriched in cardiac-like markers (Fig 4A) and is linked to ‘aortic valve and endocardial cushion morphogenesis’ (Fig S4A). In contrast, cluster 20 is largely associated with neural-like GOs (Fig S4B) and enriched in neural crest markers *HOXA3/B3* and *PDGFRB* (Fig 4A). These observations suggest two separate embryonic origins for cluster 20 and cluster 4, neural crest and secondary heart field respectively. From the set of differentially expressed genes, we largely selected TFs, which define lineage identity, to construct two distinct gene modules. Specifically, the ‘TBX20 SOX6 GATA4 PRRX1’ module is highly expressed in embryonic cluster 4, while the ‘MEIS1 JAG1 ROR1 PRDM6’ module is enriched in embryonic cluster 20. These eight developmental genes were sufficient to segregate the embryonic nuclei into two distinct subtypes (Fig. 4B), which we designated as group 1 and group 2. Group 1 consists entirely of nuclei from cluster 4, while group 2 is primarily composed of nuclei from cluster 20, with a smaller contribution from cluster 4.

**Figure 4.**
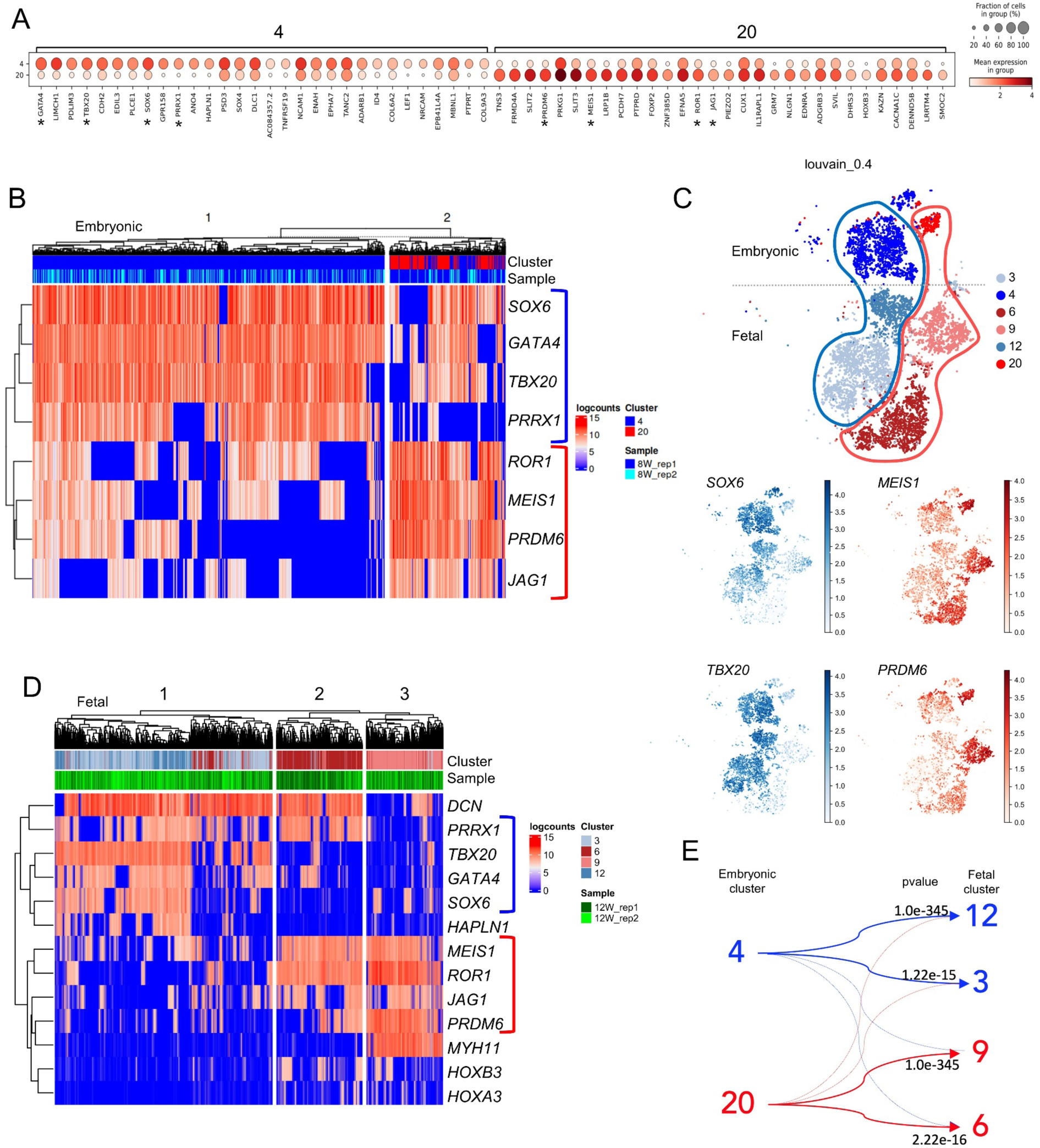
Lineage deconvolution of embryonic and fetal nuclei. A. Pairwise differential gene expression of the two embryonic mesenchymal clusters; genes chosen for gene modules are marked by asterisks. B. Heatmap using embryonic gene modules, obtained using k-means clustering, separates embryonic mesenchymal nuclei in two groups. C. Mesenchymal cell clusters of embryonic and fetal time points, projected on a two-dimensional tSNE and labelled using gene modules. Fetal clusters derive from separate ‘blue’ and ‘red’ embryonic lineages. D. Heatmap of embryonic gene modules and cell type marker genes using k-means clustering identifies three main groups of fetal nuclei. E. Lineage trajectories of embryonic and fetal nuclei. Using the entire fetal datasets, cluster 3 and 12 nuclei are identified as descendent of embryonic cluster 4, while cluster 6 and 9 are the most likely descendant of embryonic cluster 20, consistent with the use of gene modules in the mesenchymal subset of fetal nuclei in 3D.

Next, we asked if embryonic gene modules are inherited by fetal nuclei. Indeed, we found that the expression of our embryonic gene modules largely segregates developmental nuclei into two distinct lineages (Fig 4C): a ‘blue’ lineage, composed mainly of clusters 4, 3, and 12, which exhibits higher expression of module 1 (SOX6, TBX20) and a ‘red’ lineage, including clusters 20-6-9, which shows higher levels of module 2 expression (*MEIS1, PRDM6*). We then combined embryonic gene modules with markers of differentiated cell types, obtaining three distinct groups of 12pcw mesenchymal nuclei (Fig 4D). Module 1 embryonic signature is inherited by fibroblasts and valvular cells, corresponding to cluster 3 and 12, respectively (Fig 4D), suggesting a common embryonic progenitor for these cell types. Since module 1 defines the secondary heart field–derived embryonic cluster 4, we conclude that fetal group 1 nuclei derive from the secondary heart field. Conversely, the module 2 signature was inherited by two distinct cell types: smooth muscle cells (cluster 6) and fibroblasts (cluster 9). These groups were further segregated by the expression of cell type-specific markers, *MYH11* (group 2) and *DCN* (group 3). Module 2 is primarily associated with cluster 20, which is enriched in neural crest markers, with a smaller contribution from cluster 4, suggesting that fetal group 2-3 nuclei derive from both the neural crest and secondary heart field. Consistent with this, expression of cardiac neural crest markers, *HOXA3/B3,* is almost exclusively confined to ‘group 2-3’ fetal nuclei, supporting the notion that these nuclei (mainly clusters 6 and 9) partially derive from neural crest progenitors. This is in line with observations in the mouse model ^61^ where smooth muscle cells in the aortic root originate from both neural crest and secondary heart field progenitors.

To confirm the lineage relationships, we developed a robust method to trace embryonic signatures in fetal cells. We expanded the gene module repertoires to include the top 100 most distinctive genes from the mesenchymal progenitors of clusters 4 and 20 (Table S5) and examined fetal nuclei populations that retained expression of these genes. When applied to the entire 12pcw dataset (including cardiac, endothelial and mesenchymal clusters), our method confirmed the same lineage relationship between embryonic and fetal mesenchymal nuclei that were previously identified using gene modules (Fig 4E). In sum, our analysis indicates that the two spatially distinct mesenchymal populations in the fetal OFT, smooth muscle cells and valvular fibroblasts (Fig 3J-J^I^; K-K^I^) derive from separate embryonic populations: cluster 20 for smooth muscle cells and cluster 4 for valvular fibroblasts. In contrast, the *DCN*-positive, *HAPLN*-negative fibroblast cells, which spatially intermingle in OFT tissues, derive from both cluster 4 and 20 embryonic progenitors (Fig S4C).

### Cellular constituents of adult aortic valves

Semilunar (aortic and pulmonary) valves are the only distinctive and recognizable derivatives of the OFT that are retained in the adult. For adult samples, we focussed on the aortic valves because of their frequent association with disease, both genetic (bicuspid aortic valves) and adult (valve calcification) disease. We collected female samples to mitigate individual variability and maximise the possibility to analyse healthy aortic valves, justified by the lower incidence and severity of aortic disease in females versus males. Histologically normal aortic valves were procured from healthy adult hearts collected for transplantation and subsequently rejected. A total of 5,430 single nuclei from three human aortic valve samples (Fig 5A) were segregated into 11 different clusters by unsupervised clustering (Fig 5B). Leveraging established markers, we could separate the clusters into three major compartments, interstitial (7 clusters, largely *NAV2*-positive), endothelial (2 clusters, *CDH5*-positive) ^62^ and immune (2 clusters, *ITGAM*-positive macrophages and *IKFZ1*-positive dendritic cells) ^63, 64^ (Fig 5C).

**Figure 5.**
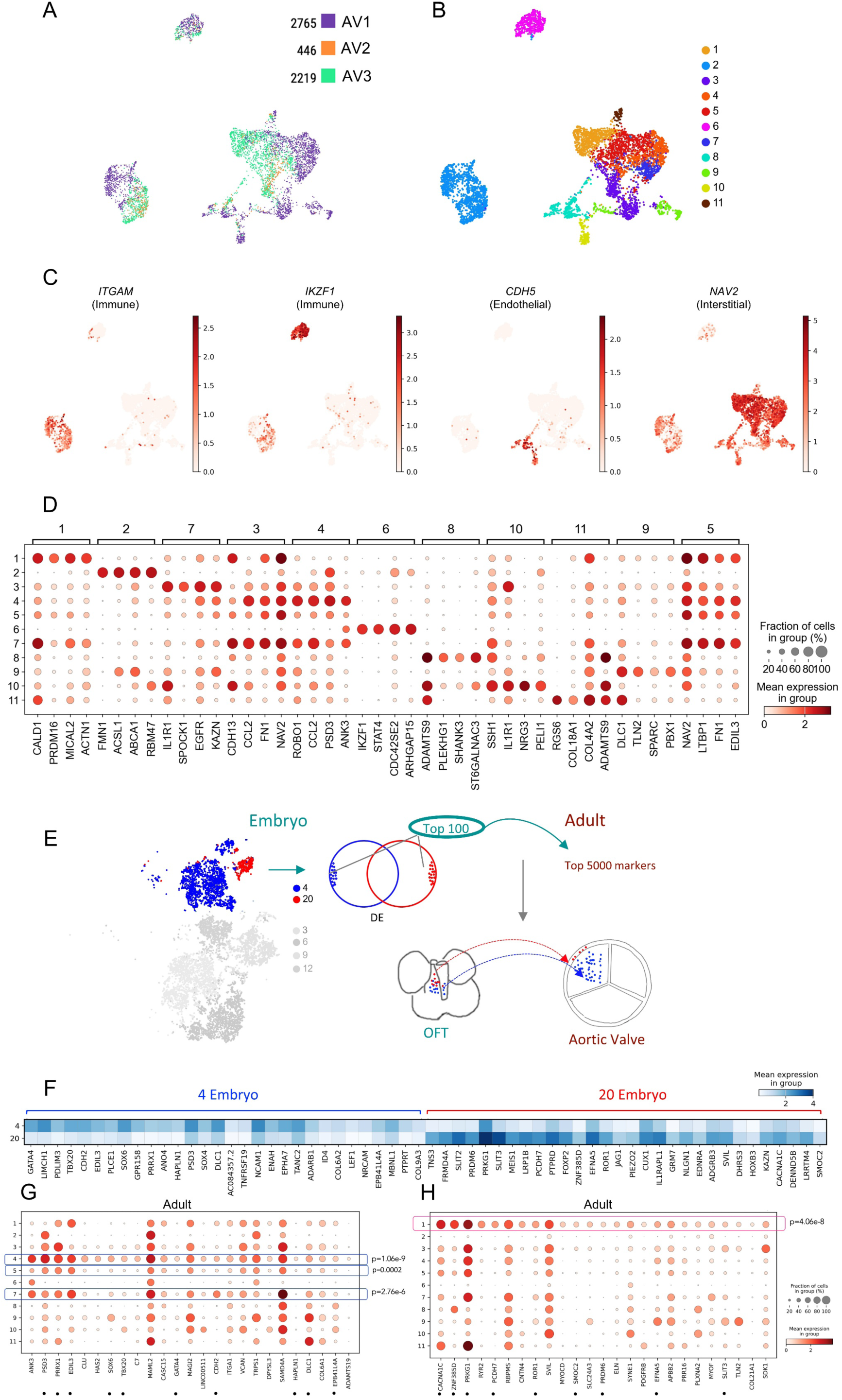
Cellular constituents of the mature aortic valves. A. Aortic valve sample association projected on a two-dimensional UMAP. Nuclei are colored by sample. B. Nuclei clusters visualized by unsupervised clustering. Nuclei are colored by cluster. C. Cell lineages identified using established lineage-specific markers. Each nucleus is colored based on the scaled expression of the indicated marker. D. Top differentially expressed genes in clusters identify known lineage markers. E. Overview of the method used to trace adult descendent of embryonic nuclei. The first step is the identification of distinctive signatures in embryonic progenitors; for this, we used the top 100 differentially expressed (DE) genes in our chosen progenitor populations, cluster 4 (blue) and cluster 20 (red). The second step is the identification of the top 5000 marker genes for each adult population; this is done by comparing each cluster with the rest of the dataset. Finally, we search for the 100 DE embryonic genes in the marker genes of adult clusters. Adult clusters with top hits are identified as the descendent of the embryonic lineage; the statistical significance is calculated using a hypergeometric test. F. Dotplot displaying the 30 top DE genes (mean expression values) in embryonic cluster 4 and cluster 20, respectively. The same dot plot, previously shown in Fig 3C, has been included here to facilitate cross-comparison with Fig 6GH. G. Distribution of cluster 4 embryonic signature genes in adult nuclei clusters. Cluster 4, 7 and 5 express a highly significant fraction of embryonic cluster 4 genes. Top 30 DE genes in embryonic cluster 4 and cluster 20 (shown in F) are highlighted by dots. H. Distribution of cluster 20 embryonic signature genes in adult nuclei clusters. Cluster 1 expresses a highly significant fraction of embryonic cluster 20 genes. Top 30 DE genes in embryonic cluster 4 and cluster 20 (shown in F) are highlighted by dots.

We performed differential gene expression analysis to aid in the classification of each cell cluster. Heatmaps (Fig S5A) group together immune (2,6) and endothelial cell types (8,10). In addition to *CDH5* ^65^, endothelial clusters are distinguished by high levels of *ADAMTS9,* which encodes for a metalloproteinase implicated in aortic valve anomalies in mouse ^66^ (Fig 5D)*. ADAMTS9* is also expressed in cluster 11, which is *CDH5*-negative (Fig 5D). Of the remaining clusters, cluster 1 is transcriptionally distinct and contains *CALD1-, ACT1*-positive SMC. AV3 is highly enriched in this cluster (26% vs 5.8% and 2.9% in AV1 and AV2 respectively) (Fig S5B); we attribute this skewed enrichment to aortic wall tissue being sampled with the aortic valves in AV1, as SMC are not a main constituent of the aortic valves, rather than to intrinsic variability across samples. Indeed, when cluster 1 is removed, the three samples are consistently similar to each other (Fig S5C). Cluster 11, which accounts for a small proportion of nuclei in all three samples (ranging from 1.5% to 2.3%) was the most related cluster to cluster 1 (Fig S5B; Fig 5E).

Clusters 3,4,5,7 display expression of similar transcripts, such as *NAV2*, *LTBP1*, *FN1* and were assigned to valvular interstitial cells (VIC) (Fig S5A; Fig 5D). VICs deposit three highly organized layers (fibrosa, spongiosa and ventricularis) of extracellular matrix (ECM), that compose the mature valve structure ^67^. These cells are identified by the expression of transcripts encoding for ECM proteins, most notably *COL1A1*, *VIM*, *COL3A1*, *VCAN*, *BGN* and *LUM*. ECM encoding transcripts were enriched in clusters 3,4,5,7 (VIC) across all the adult samples examined (Fig S5D), with *VIM* and *VCAN* displaying a broader expression. Differently from cluster 5, clusters 4 and 7 express high levels of *CCL2*, an inflammatory cytokine ^68^, suggesting nuclei in these clusters correspond to activated fibroblasts. Finally, cluster 9 displays enrichment in *DLC1* and *TLN2*, implicated in cytoskeletal changes ^69, 70^, and *SPARC* (osteonectin), implicated in valve calcification ^71^. While the above clusters are represented across all samples, AV1 clusters 4-7-9 contain on average more nuclei relative to AV2 and AV3 (Fig 5D; Fig S5E). The increased expression of *SPARC* (cluster 9), combined with higher levels of *CCL2* (cluster 4 and 7) in AV1, indicates the possibility of inflammatory processes that could eventually lead to valve calcification in AV1 sample. We did not detect myocardial cells in the adult valve tissue, consistent with evidence that myocardium contributes to early arterial root and cushion formation but does not persist in mature valves. Myocardial gene expression is already absent from the valve leaflet cluster by CS16-19^22^. Our adult dataset therefore reflects the valve complex and adjacent arterial root region, a subset of embryonic OFT derivatives rather than the entire OFT myocardium.

### Persistence of embryonic signatures in adult cells

Whilst differentiated, 12pcw nuclei maintain robust embryonic signatures (Fig 4DE). We asked if such signatures also persist in terminally differentiated adult cells and could be used as labels to retrospectively attribute cellular relationships between embryo and adult cells. To address this, we leveraged our method to identify embryonic signatures in adult cells. Using the top one hundred most distinctive genes of the embryonic mesenchymal progenitors (cluster 4 and 20 in the embryo) (Table S5), we looked for populations of adult nuclei enriched in the expression of most of these genes (Fig 5E). We found that adult clusters 1,4,7 retain expression of a highly significant fraction of distinctive embryonic genes, with clusters 4,7 largely maintaining embryonic cluster 4 expression signature (blue) (Fig 5FG), while cluster 1 displays a significant match with embryonic cluster 20 (red) (Fig 5FH). The finding that smooth muscle cells (adult cluster 1) are derived from embryonic cluster 20 is consistent with cluster 20 being the source of smooth muscle cells at 12pcw (Fig 4). Valvular fibroblasts (clusters 4,7) derive from the embryonic population in cluster 4, which is also linked to fetal valvular fibroblasts (Fig 4). This result is significant because our embryonic signatures derive from undifferentiated cells which are yet to express obvious differentiation markers, suggesting that adult cells retain their ancestral embryonic make up. The expression of a representative gene set from the 100-gene embryonic signatures was projected onto adult valve cells, confirming the findings shown in Fig 5F-H (Fig S7AB). As our adult clusters reflect aggregation of three individual samples, we asked if distinctive embryonic signatures can be detected despite confounding individual factors (ageing, environmental exposure, genetic background, etc). We performed the same analysis on individual samples AV1 and AV3 (AV2 was excluded due to the lower number of nuclei). We independently re-clustered each sample and applied our method to identify the descendent nuclei of embryonic cluster 4 within individual samples. Our method linked embryonic cluster 4 to cluster 0 in AV1, and to cluster 2 and 3 nuclei in AV3 (Fig S6AB); these clusters contain the majority of adult cluster 4 aggregate nuclei (Fig S6CD). This indicates that distinctive embryonic signatures can be detected over individual variability. In sum, our analysis indicates that distinctive patterns of embryonic gene expression persist in adult cells, can be consistently detected in individual adult samples, and can be used as labels to retrospectively attribute cellular relationships between embryo and adult cells. Finally, as the level of expression of a gene across different cells can provide an initial indication of its functional role, we explored the relative expression levels of distinctive embryonic signatures in embryonic nuclei and their adult descendent nuclei. Generally embryonic genes display lower expression levels in adult cells relative to their embryonic progenitors (Fig S6E-F).

### Spatial profiling of OFT defect genes reveals candidates for congenital heart malformations

OFT defects occur when the vessels leaving the heart do not form or remodel correctly, resulting in problems with blood circulation and/or oxygenation in the neonate ^72^. The most severe include common arterial trunk (CAT), transposition of the great arteries (TGA), double-outlet right ventricle (DORV) and Tetralogy of Fallot (TOF). Using spatial transcriptomics, we investigated the distribution of genes whose mutations cause OFT defects (Fig 6A-A^I^). *JAG1* mutations cause TOF ^73^; *JAG1* transcripts largely concentrate around the aorta, in the semilunar valves and the septum (Fig 6B-B^I^). Mutations in *GATA* family members affects OFT formation in different ways, leading to DORV (*GATA5*) ^74^, TOF (*GATA4, GATA6*) ^75, 76^ and CAT (*GATA6*) ^43^. Different from *GATA5*, which shows a restricted expression to the vessels (Fig 6C, C^I^), the distribution of *GATA6* and *GATA4* transcripts is broader. *GATA6* is detected in the vessels, the septum and cardiac tissue (Fig S8A), while *GATA4* transcripts are more prominent in cardiac tissues (Fig S8B). Mutations in *GATA4-5-6* also causes BAV, a less severe condition characterized by two valves instead of three ^77^. Similar to *GATA5*, *GATA6* is highly expressed in the valves, marked by *HAPLN1* (Fig 3K^I^). As a final example, mutations in *NR2F2* cause DORV and TOF ^78^. Differently from the previous examples, *NR2F2* transcripts are largely excluded from OFT structures and are mainly located in cardiac tissues surrounding the aorta and the pulmonary artery (Fig 6D, D^I^).

**Figure 6.**
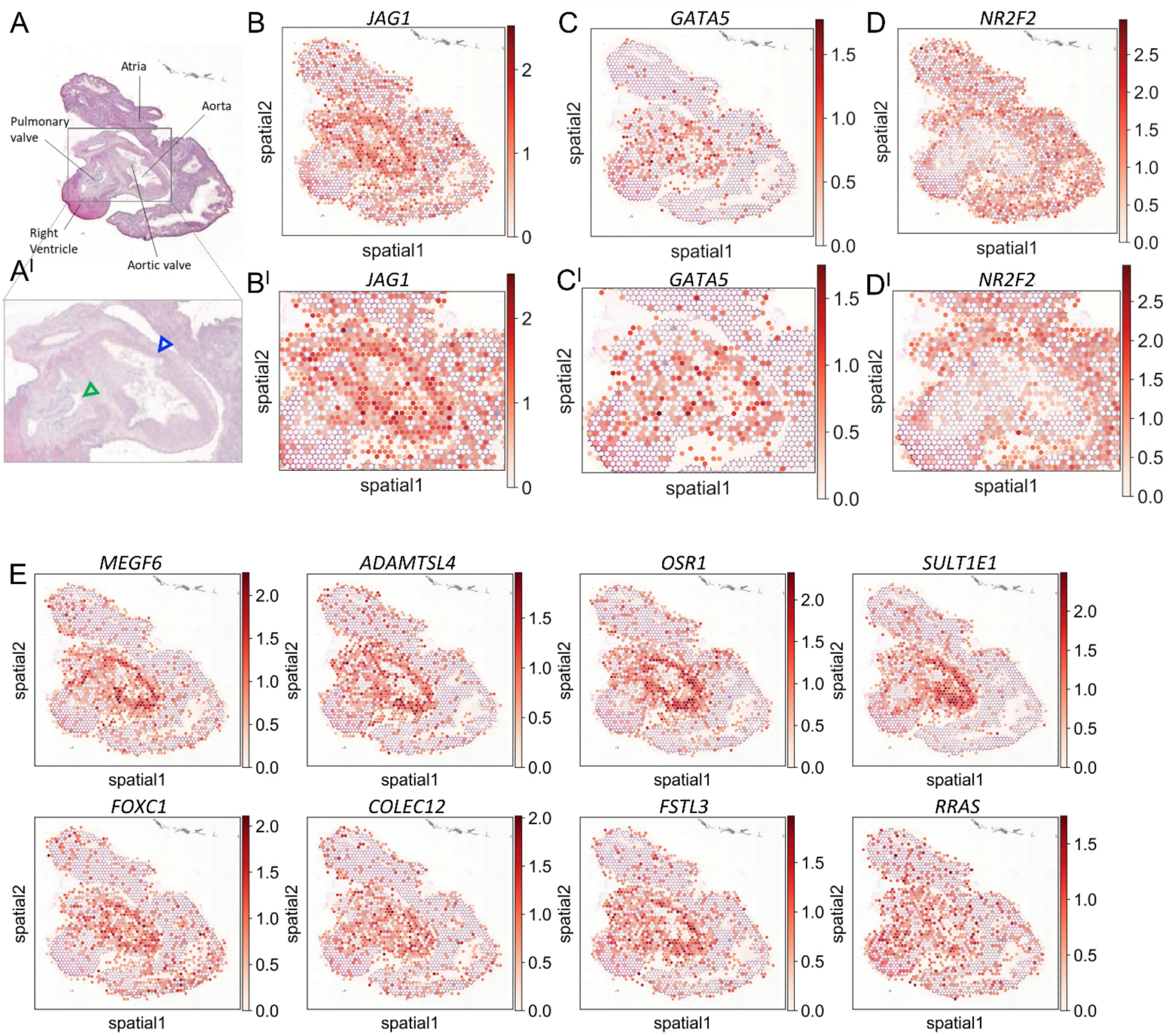
Genes mutated in congenital OFT defects. A. H&E staining of spatial transcriptomics section (Fig 2Jc), and magnified view of aortic and pulmonary valve area (A^I^). The aorta and pulmonary trunk are indicated by blue and green arrowheads, respectively. B-D. *JAG1* (B), *GATA5* (C) and *NR2F2* (D) gene expression patterns on the same section. B^I^- DI, Genes as in BD with corresponding magnification of valve area (B^I^, C^I^, D^I^). E. Genes identified as displaying spatially similar expression pattern to *JAG1*.

We reasoned that genes exhibiting a distribution pattern similar to, or matching those mutated in OFT defects, could serve as potential candidates for OFT anomalies. To test this hypothesis, we focused on *JAG1*, whose transcripts exhibit a distinct pattern around the vessels, and generated an algorithm to identify genes that display comparable expression patterns to *JAG1* (Fig 6E). The algorithm filters out any gene that is not expressed in the majority of *JAG1*-positive spots (>50%) and is also expressed in *JAG1*-negative spots (see methods). Using this approach, we identified *FOXC1* and *OSR1,* whose loss of function in mouse results in semilunar valve abnormalities and aortic arch coarctation, and defects in heart septation, respectively ^79, 80^. In addition, variants in *FOXC1* were recently identified in patients with conotruncal heart defects ^10^.

## Discussion

We used snRNA-seq to analyze 30,166 nuclei, which covered two stages of OFT development (CS16-17 and 12pcw) up to the adult derivatives of the OFT, the aortic valves. In parallel, we used spatial transcriptomics to define the distribution of fetal cell populations and visualize the expression patterns of genes implicated in OFT defects, which make up one third of all cases of CHD. These datasets provide a large reference framework of OFT cell repertoires and their gene expression profiles, and constitute a valuable resource for enhancing our understanding of OFT defects and advancing the discovery of previously unknown genes implicated in these conditions. A major finding emerging from our analysis of a timeline of human development is that adult cells retain their ancestral embryonic signatures, which may have implications for adult-onset aortic valve disease.

During human embryo development, the OFT undergoes formation and remodelling between CS13 and CS23. This process results in the separation of the aorta and pulmonary trunks, creating the double circulation found in mammals. It also leads to the formation of the arterial valves, which continue to mature after the arterial vessels have separated. Focussing primarily on mesenchymal cells, which are responsible for building the semilunar valves and the separation of the aorta from the pulmonary artery, we identify two distinct groups of embryonic progenitors at the earliest stage (CS16/17). These mesenchymal nuclei do not yet express clear differentiation markers, but display distinctive neural crest and secondary heart field expression signatures. In contrast, by 12pcw mesenchymal nuclei express cell type markers and cluster into four subgroups, corresponding to fibroblast-like cells, smooth muscle cells, valvular interstitial cells, each localized to distinct regions within the OFT. Using gene modules and a new lineage tracing tool, we traced these four fetal mesenchymal populations back to their embryonic precursors. Our results reveal that valvular fibroblasts and smooth muscle cells have separate embryonic origins. Valvular fibroblasts, the mesenchymal cells involved in constructing the semilunar valves, appear to originate from endothelial cells. This aligns with the finding that mouse arterial valves derive from endothelial cells undergoing EMT in the endocardium - the myocardium-adjacent cell layer ^55, 56^. The mesenchymal progenitors of fetal valvular fibroblasts and adult VICs are regulated by a GATA6-driven gene network. Mutations in *GATA6* are a known cause of BAV, suggesting that this regulatory program plays a critical role in semilunar valve development. The GATA6 regulon includes genes whose mutations are associated with OFT and valve defects in mice, as well as aortic valve disease in humans. Although we did not uncover a novel single-gene marker specific to humans (analogous to LRG5^13^), our identification of a GATA6 network highlights molecular mechanisms downstream of GATA6 that drive valve formation, advancing our understanding of normal valve development and informing the search for genes involved in BAV susceptibility and aortic valve disease. In contrast, smooth muscle cells derive from a distinct CS16-17 embryonic population. A consistent set of these cells displays expression of cardiac neural crest markers, in agreement with previous observations in mice, indicating that smooth muscle cells in the human aortic root originate from the neural crest and secondary heart field ^81^. Finally, both embryonic mesenchymal populations also give rise to non-valvular fibroblast cells (*DCN*^+^, *HAPLN1^-^)*.

We find that distinctive embryonic signatures persist into adulthood. While previous studies have described that reactivation of fetal programs in adult heart disease ^82, 83^, all our adult samples are derived from healthy individuals. This suggests that the persistence of early developmental signatures is a pervasive feature of normal adult cells, not merely a pathological response. Single-cell analyses have revealed high levels of heterogeneity within the same cell type. Our findings suggest that the inheritance of ’embryonic memories’ by adult cells may be a major contributing factor to this heterogeneity. In support of this, organ-specific developmental signatures have been observed in adult fibroblasts ^84^. In the human context, the persistence of distinct embryonic signatures in adult cell types may influence their susceptibility to disease, and potentially contribute to a better understanding of disease heterogeneity.

Connecting cell lineages over extended periods of time can be challenging. Existing tools typically use single-cell RNA sequencing to capture the complete expression profiles of individual cells in a single experiment, treating each cell as a unique time point on a continuum ^60^. However, the significant transcriptional changes that occur as cells transition from embryonic to fetal and adult cell types pose a challenge when comparing and linking cells over prolonged periods based on their global expression profiles. The persistence of embryonic signatures into adulthood opens the possibility of using these enduring molecular "labels" to trace developmental ancestry in complex tissues, particularly in contexts where related cells are temporally distant or phenotypically divergent, such as in aging or cancer.

The observation that embryonic expression signatures persist in adult cells raise obvious questions about functional significance. We find that embryonic genes are generally expressed at lower levels in adult cells relative to embryonic cells (where they are known to have a function); their persistent expression may reflect fortuitous remnants of developmental histories. Alternatively, these retained embryonic gene expressions could serve as preserved developmental blueprints, ready to be swiftly and efficiently reactivated when the need arises, such as during injury or tissue repair. Reinforcing this perspective, heart failure is associated with the reawakening of a fetal gene program ^82, 83^.

In summary, our work extends beyond confirming previously reported cell types by (i) defining a GATA6-regulated human valve progenitor lineage and its derivatives, (ii) establishing distinct embryonic origins for smooth muscle and valvular fibroblasts, and (iii) demonstrating the persistence of embryonic signatures in adult valve cell populations. These conclusions are directly supported in tissue by our spatial transcriptomics data, which map these lineages and regulatory programs to defined anatomical domains within the human OFT and semilunar valves.

## Supporting information

Supplentary Figures_Legends for Supplementary figures and Tables

## Acknowledgements

We thank Andy Hayes and the other members of the Genomic Technologies Core Facility, Roger Meadows of the Bioimaging facility and Gareth Howell of the Flow Sorting Core Facility at the University of Manchester. We also thank Zoulfia Darieva, Peyman Zarrineh, Rachel Jennings and Aoibheann Mullan for help and discussions. A special thanks to Jasmin Turner for help with cryosectioning and histology. This work was supported by joint funding from the Medical Research Council (MRC) (http:www.mrc.ukri.org) grant MR/S03613X/1 and British Heart Foundation (BHF) grant SP/18/12/34300 to NB and SDB. The Human Developmental Biology Resource (www.hdbr.org) is funded jointly by the Medical Research Council and the Wellcome Trust (MR/R006237/1). Adult tissue acquisition was funded by the National Institute for Health and Care Research (NIHR) Blood and Transplant Research Unit in Organ Donation and Transplantation (NIHR203332), a partnership between NHS Blood and Transplant, University of Cambridge and Newcastle University.

## Data availability

All data described have been deposited in ArrayExpress with accession numbers listed in table S1 and will also be made available upon acceptance of the manuscript through the Human Cell Atlas. The spatial transcriptomics data is accessible at https://spatialtranscriptomics-uom.bmh.manchester.ac.uk/. Instructions on how to use the app are available at the cellxgene VIP website: https://interactivereport.github.io/cellxgene_VIP/tutorial/docs/how-to-use-cellxgene-vip.html

## Code availability

The code used in Fig. 5 will be made publicly available through GitHub upon publication.

