## Supplementary material for "A cell atlas of the developing human outflow tract of the heart and its adult aortic valve derivatives": Supplentary Figures_Legends for Supplementary figures and Tables

### Figure S1

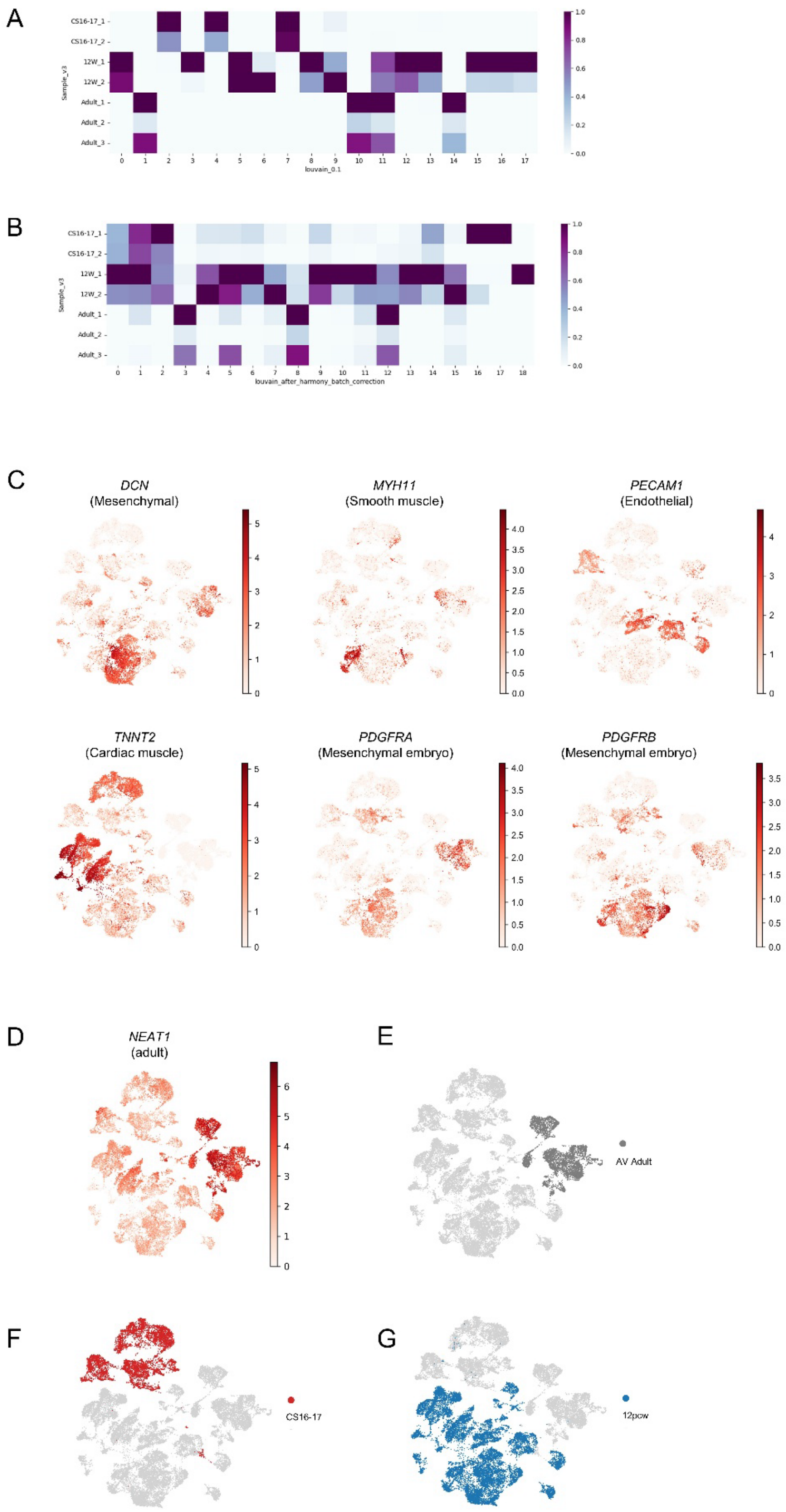
A
CS16-17_1
CS16-17_2
12W_1
12W_2
Adult_1
Adult_2
Adult_3
Sample_v3
louvain_0.1
B
louvain_after_harmony_batch_correction
C
DCN
(Mesenchymal)
MYH11
(Smooth muscle)
PECAM1
(Endothelial)
TNNT2
(Cardiac muscle)
PDGFRA
(Mesenchymal embryo)
PDGFRB
(Mesenchymal embryo)
D
NEAT1
(adult)
E
AV Adult
F
CS16-17
G
12pcw

**Figure S1**. AB. Confusion matrix of Louvain clusters' distribution across samples before (A) and after (B) Harmony batch correction. Values (number of nuclei in each cluster) are scaled by min max scaling. A. Clusters are composed of time-point matched biological replicates, showing that our datasets are not impacted by technical batch effects and that the variability we observe reflects meaningful changes across time. The only exception is cluster 11 (immune): immune cells are relatively homogenous across populations, and their assignments to different samples across two time points, fetal and adult, indicates batch effect is not creating undue artefacts in our data integration. B. Most clusters include samples from different time points, which indicates a loss of biological variability across time after batch correction. C. Cell types in Fig 1E labelled using established lineage markers. *DCN* identifies fibroblast populations, *MYH11* labels SMCs, *PECAM1* is a marker for endothelial cells, *TNNT2* is a marker for cardiac cells, and *PDGFRA* and *PDGFRB* mark mesenchymal cells before differentiation. Each nucleus is colored based on the scaled expression of the indicated marker. D. *NEAT1* expression. E-G. UMAP visualization of samples aggregated by stage: adult aortic valves (E, dark grey), embryonic (F, red), fetal (G, blue).

### Figure S2

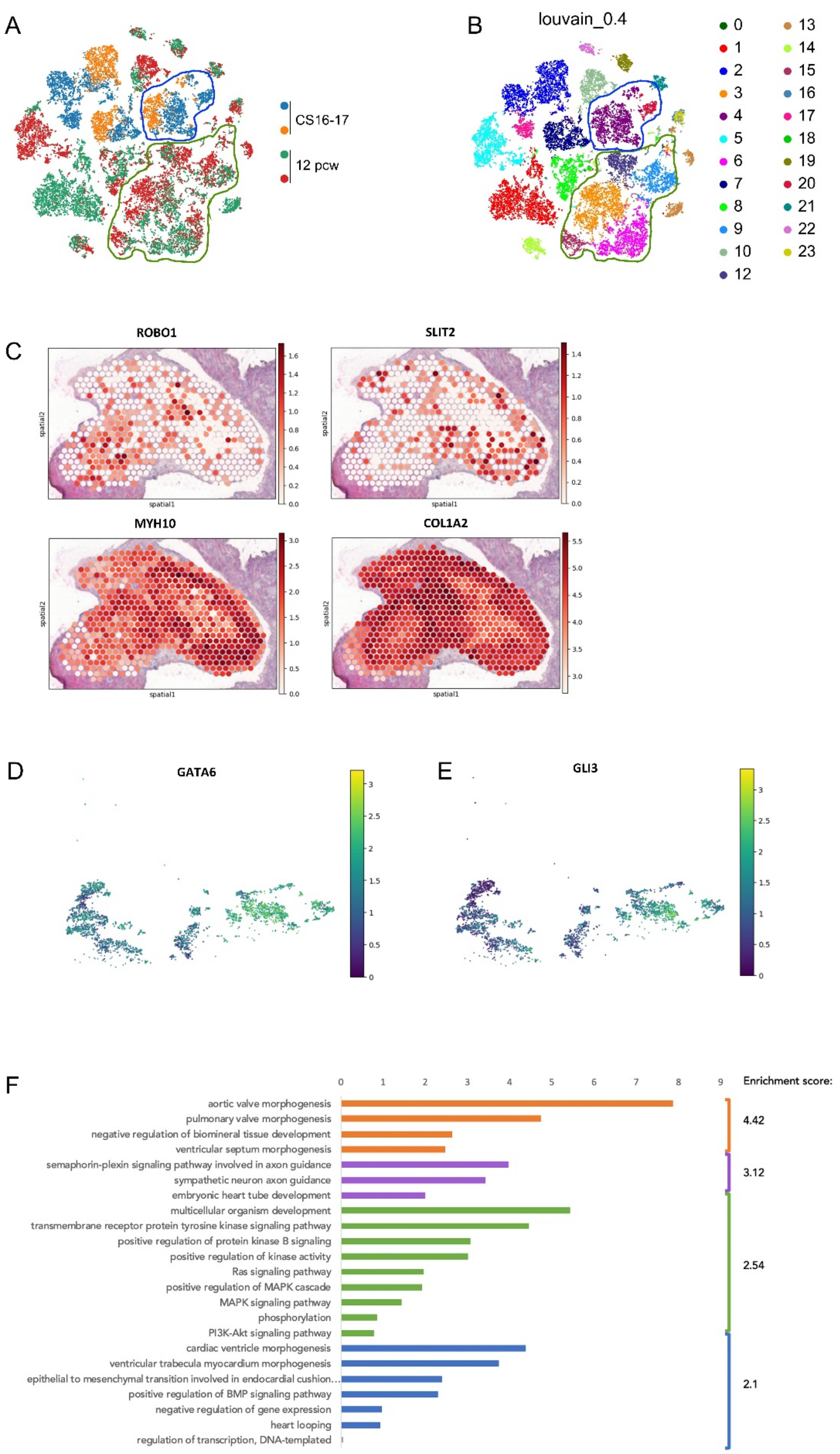

A
CS16-17
12 pcw
B
louvain_0.4
0 1 2 3 4 5 6 7 8 9 10 12 13 14 15 16 17 18 19 20 21 22 23
C
ROBO1
SLIT2
MYH10
COL1A2
spatial1
spatial2
D
GATA6
E
GLI3
F
Enrichment score:
aortic valve morphogenesis
pulmonary valve morphogenesis
negative regulation of biomineral tissue development
ventricular septum morphogenesis
semaphorin-plexin signaling pathway involved in axon guidance
sympathetic neuron axon guidance
embryonic heart tube development
multicellular organism development
transmembrane receptor protein tyrosine kinase signaling pathway
positive regulation of protein kinase B signaling
positive regulation of kinase activity
Ras signaling pathway
positive regulation of MAPK cascade
MAPK signaling pathway
phosphorylation
PI3K-Akt signaling pathway
cardiac ventricle morphogenesis
ventricular trabecula myocardium morphogenesis
epithelial to mesenchymal transition involved in endocardial cushion…
positive regulation of BMP signaling pathway
negative regulation of gene expression
heart looping
regulation of transcription, DNA-templated
4.42
3.12
2.54
2.1

**Figure S2.** AB. Louvain 0.4 re-clustering of embryonic and fetal samples (without adult samples), presented by tSNE. Nuclei are colored by sample (A) and cluster (B). Embryonic and fetal mesenchymal clusters are highlighted by blue and green contours, respectively. C. Spatial Transcriptomics of Aorta (Ao) and Pulmonary Artery (PA) (clockwise): *ROBO1*, *SLIT2*, *MYH10*, *COL1A2*. D. Expression of *GATA6* (Log normalised values) in CS16-17 nuclei, projected onto the RNA velocity plot shown in Figure 2K. E. Expression of *GLI3* (Log normalised values) in CS16-17 nuclei, projected onto the RNA velocity plot shown in Figure 2K. F. Gene ontologies associated with distinctive markers of embryonic endothelial cells (cluster 7). Functional annotation clustering of top 200 genes enriched in embryonic endothelial cluster 7 (relative to fetal endothelial cluster 9, 13) was performed using DAVID and -Log10(Pv) was plotted in Excel. Genes are listed in Table S3.

Figure S3

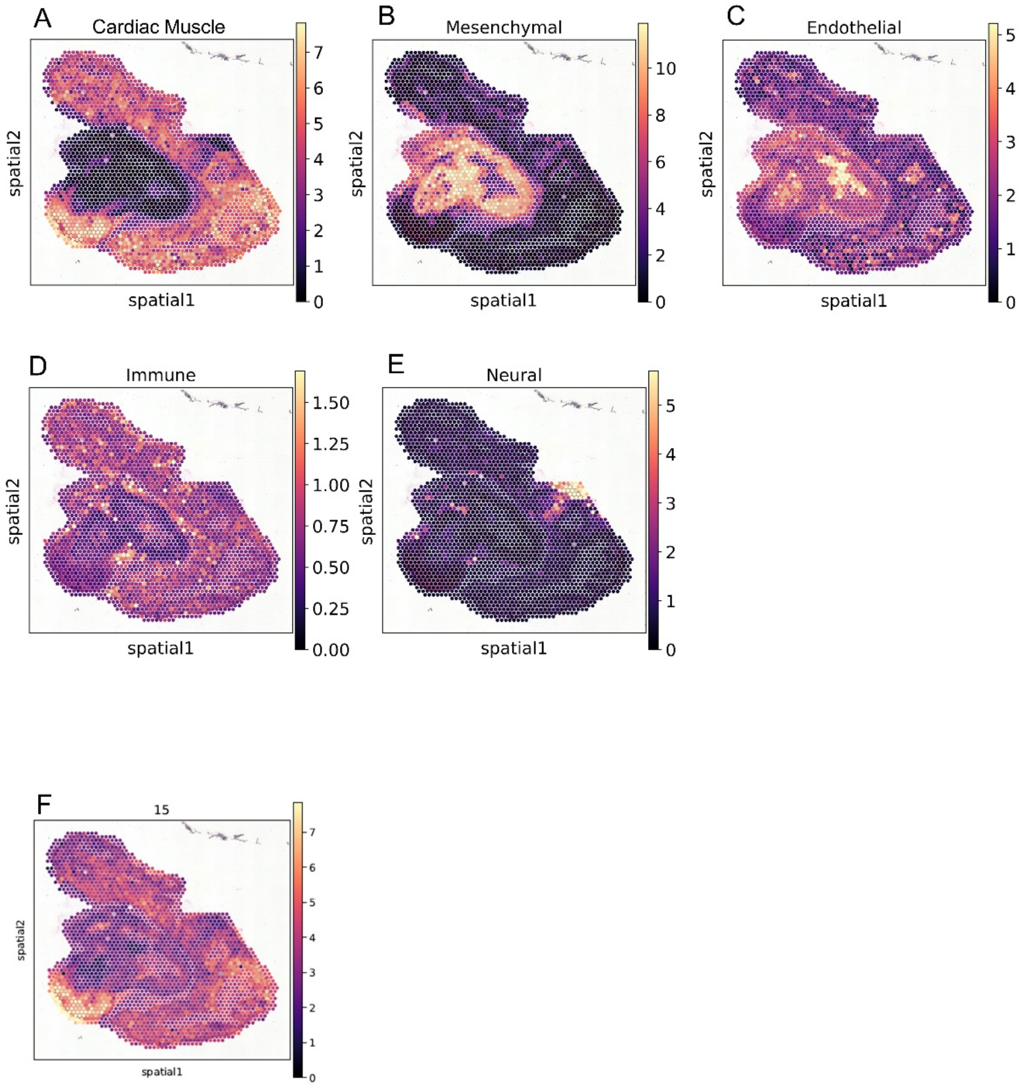


**Figure S3**. Mapping of cell types from transcriptomics data to spatial location using Cell2location. A. Cardiac muscle cells in the atria and ventricle. B. Mesenchymal cells consisting of smooth muscle cells inside the lumen of the aorta, fibroblast layer around the vessels and in the aortic and pulmonary semilunar valves. C. Endothelial cells in the aortic semilunar valves. D. Immune cells. E. Neuronal cells. F. Cluster 15 is restricted to the right ventricle and was removed from the analysis of OFT mesenchymal clusters.

### Figure S4

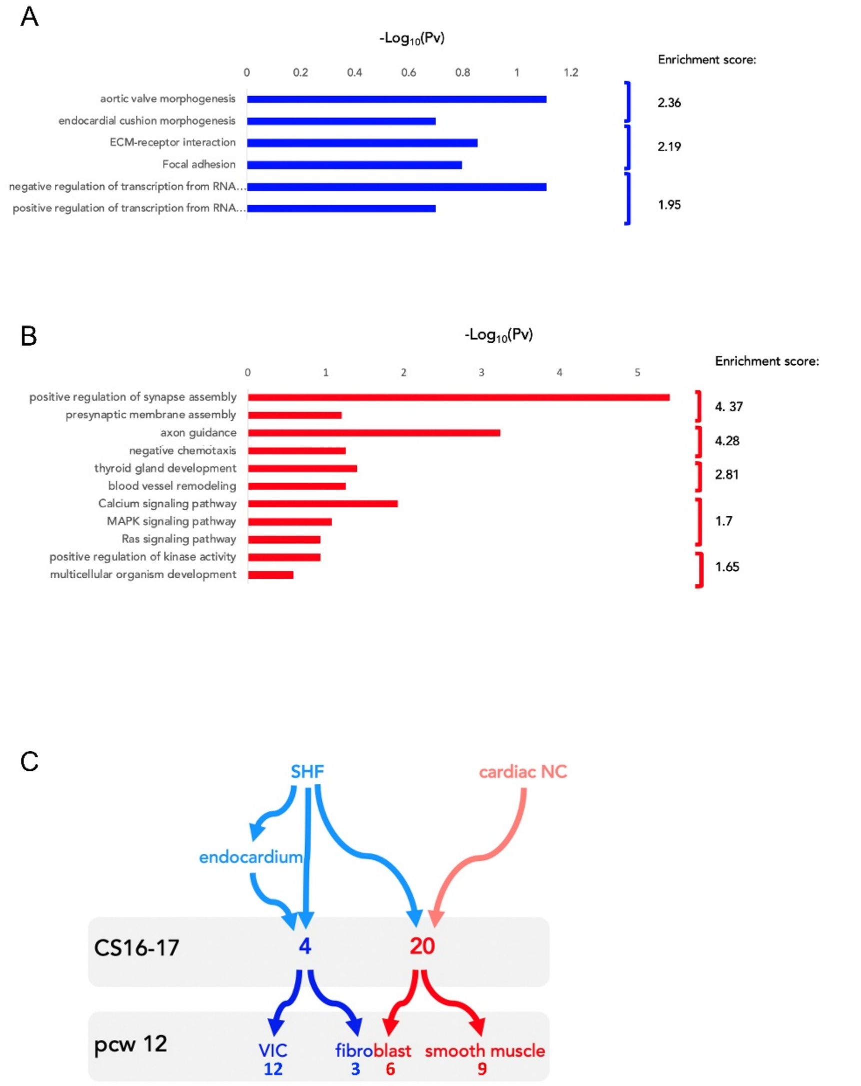


**Figure S4**. Gene ontologies associated with distinctive developmental signatures of mesenchymal subtypes. Functional annotation clustering of top 100 genes enriched in embryonic cluster 4 (A) and cluster 20 (B) was performed using DAVID and -Log10(Pv) was plotted in Excel. Cluster 4 (A) is enriched in cardiac-like markers and cluster 20 (B) in neural crest markers. Genes are listed in Table S4. C. Lineage relationships between embryonic and differentiated fetal mesenchymal cells. Cluster 4 derives from the secondary heart field (SHF) with a contribution of SHF-derived endocardial cells. It gives rise to arterial valves and fibroblasts. Cluster 4 derives from SHF and cardiac neural crest; cells in cluster 4 are the progenitors of smooth muscle cells and fibroblasts.

### Figure S5

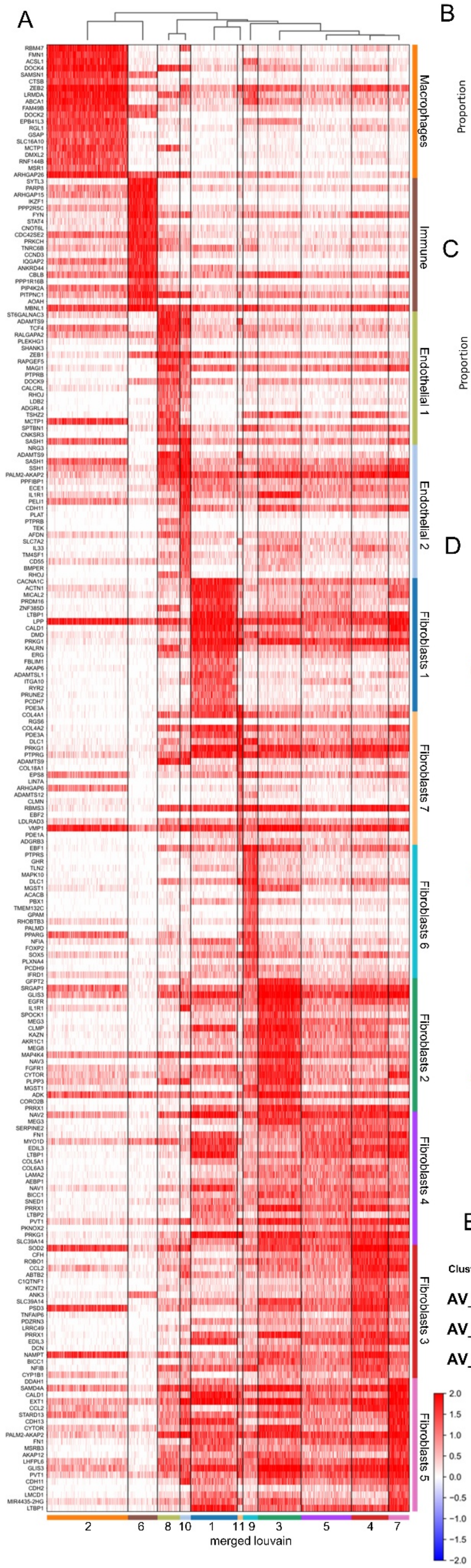


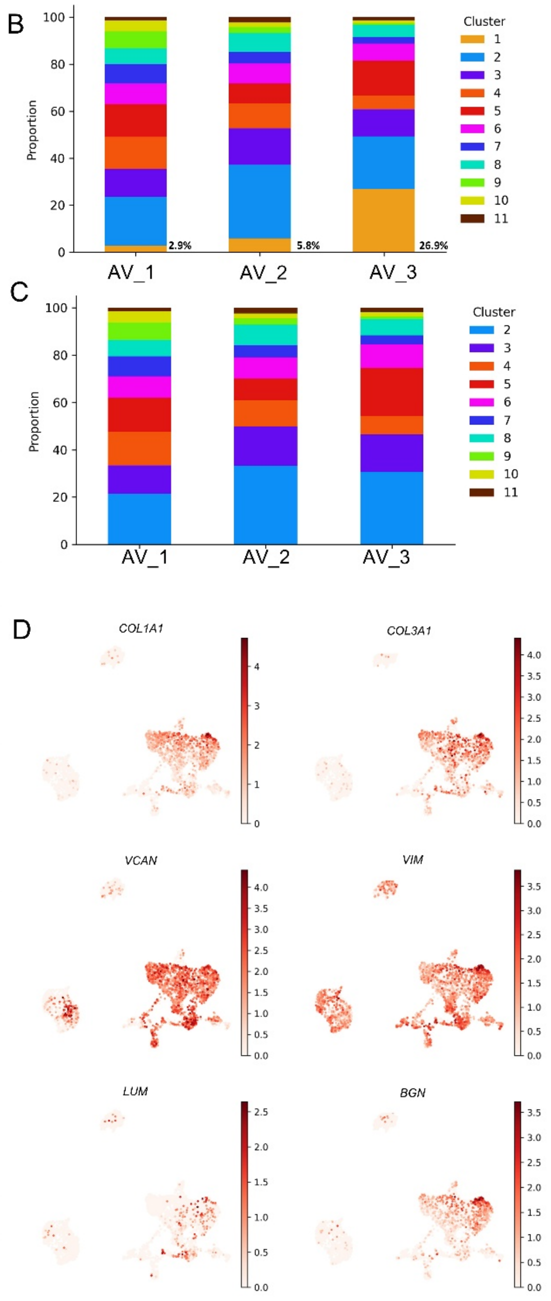


E

**clusters**

| Cluster | 2 | 3 | 4 | 5 | 6 | 7 | 8 | 9 | 10 | 11 |
|---|---|---|---|---|---|---|---|---|---|---|
| **AV_1** | 21.42 | 12.01 | 14.17 | 14.36 | 9.00 | 8.51 | 6.95 | 7.32 | 4.68 | 1.52 |
| **AV_2** | 33.33 | 16.43 | 11.19 | 9.04 | 9.04 | 5.23 | 8.67 | 2.61 | 2.14 | 2.38 |
| **AV_3** | 30.70 | 15.78 | 7.82 | 20.28 | 10.04 | 3.69 | 6.96 | 1.10 | 1.72 | 1.85 |

**Figure S5.** A. Differential gene expression heatmap of adult clusters. BC. Cluster composition in each adult sample, presented as percentage of nuclei with (A) and without (B) cluster 1. D. ECM encoding transcripts in interstitial cells. Each nucleus is colored based on the scaled expression of the indicated marker. E. Proportions of nuclei in each adult cluster.

### Figure S6

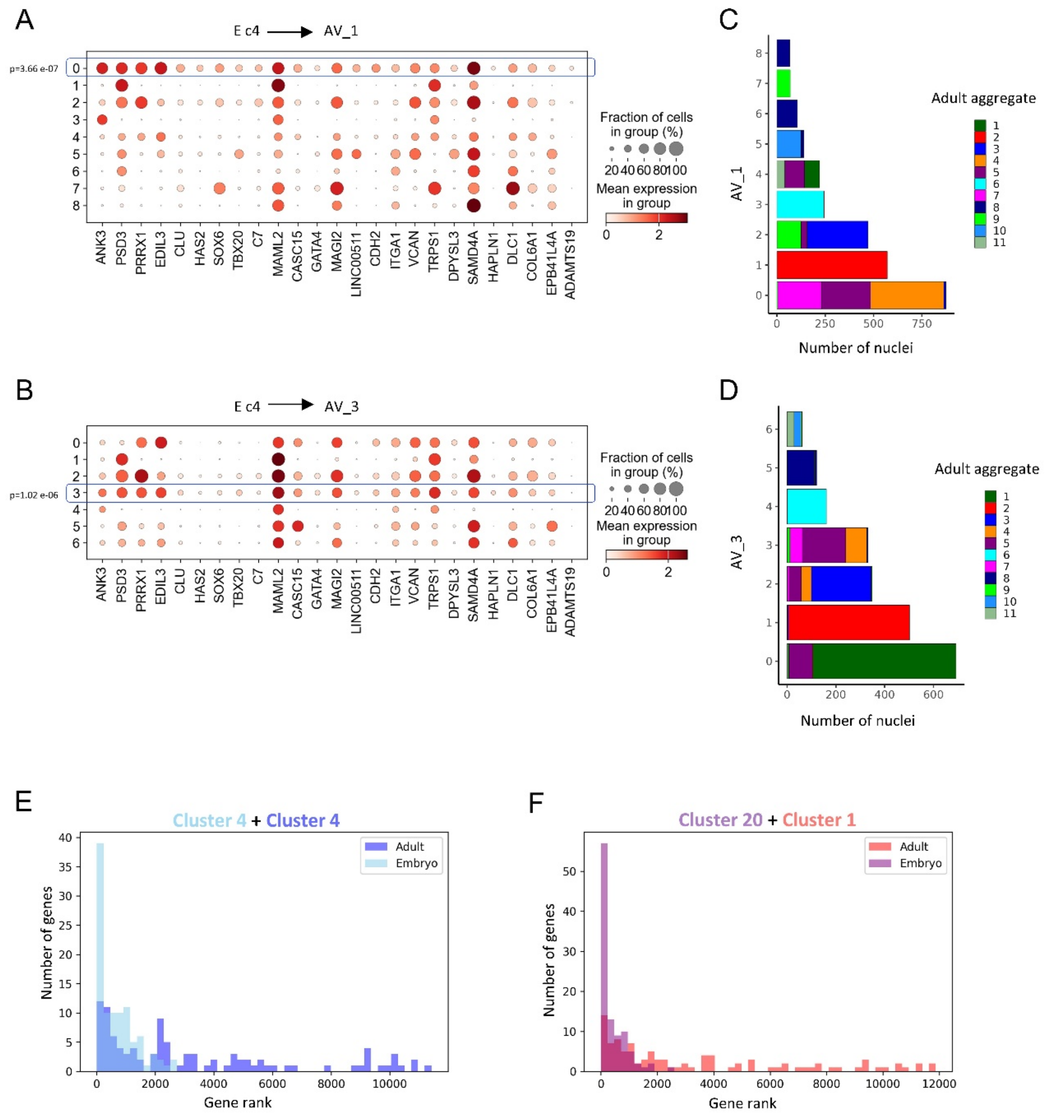


**Figure S6.** AB. Dotplots showing the distribution of cluster 4 embryonic signature genes in individual samples, AV1 and AV3. Cluster 0 in AV1 (A) and cluster 3 in AV3 (B) express a highly significant fraction of embryonic cluster 4 genes. CD. Cluster correlation between adult aggregate nuclei and individual sample nuclei. AV1 nuclei in cluster 0 contain nuclei from aggregate cluster 4,7. Similarly, AV3 cluster 3 nuclei contain the majority of aggregate cluster 4,7 nuclei. EF. Relative expression of embryonic signature genes in embryonic (E, 4; F, 20) and adult clusters (E, 4; F, 1), which are lineage related. Genes were ranked by expression values, relative to all the genes expressed in each cluster. In descendent adult nuclei, most embryonic genes (65% in adult cluster 4) are not represented in the top 1000 expressed genes. In contrast embryonic genes, are among the top 1000 expressed genes in embryonic nuclei (73% in embryo cluster 4).

Figure S7

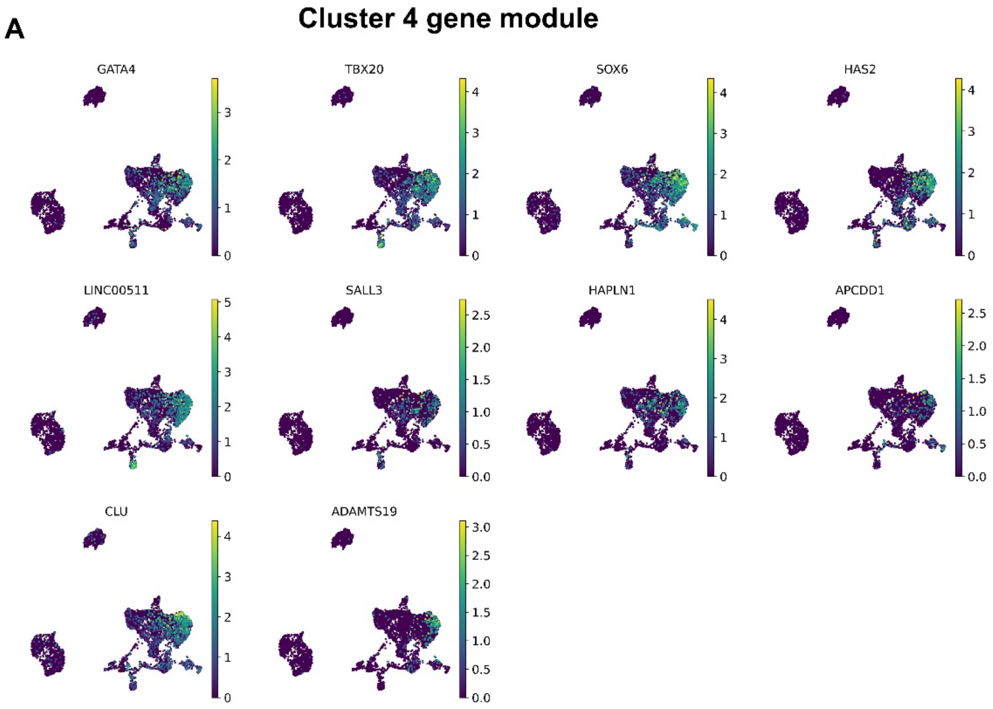


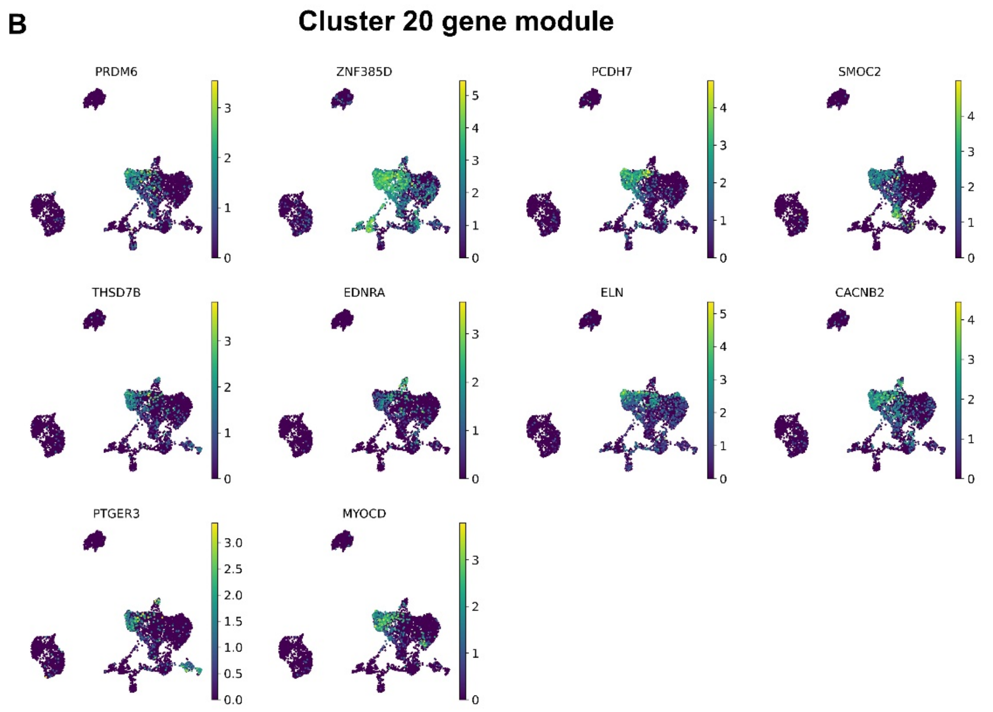


**Figure S7.** AB. Expression (Log normalised values) of 10 representative genes from the 100-gene embryonic signature lists of cluster 4 (A) and cluster 20 (B) projected onto the t-SNE shown in Figure 4E.

Figure S8

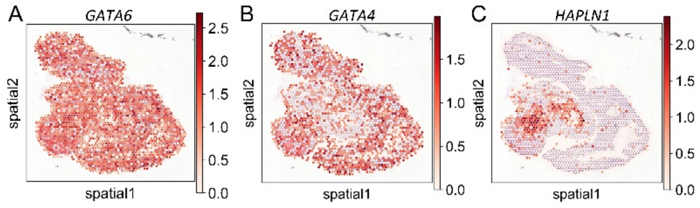


**Figure S8.** A. GATA6 and B. GATA4. GATA4 and GATA6 are broadly distributed across the OFT and surrounding cardiac tissue. Mutations in both genes cause BAV; the position of the valves is highlighted by HAPLN1 (C).

**Table S1.** Accession numbers of single cell experiments and spatial transcriptomics

**Table S2**. Marker genes (top 100) for each of the 18 clusters (0-17) in Fig1.

**Table S3**. GATA6 regulon genes associated with GATA6 binding in the OFT and pharyngeal arches in E11.5 mouse embryos. Association was established using GATA6 peaks with a fold enrichment cutoff > 10 and GREAT standard association rules.

**Table S4**. Genes enriched in cluster 7 (embryonic endothelial) relative to clusters 9-13 (fetal endothelial).

**Table S5**. Embryonic cluster 4 and Cluster 20 top 100 differentially expressed genes
